# Multi-omic profiling of human thymic B cells reveals intrathymic Ig-class switching and differentiation into multiple memory B cell subsets

**DOI:** 10.64898/2026.05.21.725517

**Authors:** Martijn Cordes, Erik B. van den Akker, Janine E. Melsen, Saskia Bunschuh, Aline Zbinden, Sandra de Bruin-Versteeg, Nannan Guo, Frits Koning, Szymon M. Kiełbasa, Karin Pike-Overzet, Mirjam van der Burg, Marcel J.T. Reinders, Frank J.T. Staal, Kirsten Canté-Barrett

**Affiliations:** Department of Immunology, Leiden University Medical Center, Leiden, The Netherlands; Leiden Computational Biology Center, Leiden University Medical Center, Leiden, The Netherlands; The Delft Bioinformatics Lab, Delft University of Technology, Delft, The Netherlands; Molecular Epidemiology, Leiden University Medical Center, Leiden, The Netherlands; Department of Biomedical Data Sciences, Leiden University Medical Center, Leiden, The Netherlands; Novo Nordisk Foundation Center for Stem Cell Medicine (reNEW), Leiden University Medical Center, Netherlands; Department of Pediatrics, Leiden University Medical Center, Leiden, Netherlands

## Abstract

Primarily recognized as the site for T cell development, the thymus supports a complex interplay between thymic stromal cells and developing thymocytes, which is essential for T cell maturation and the establishment of central tolerance. Emerging evidence indicates that thymic B cells contribute to tolerance induction by functioning as antigen-presenting cells. However, their developmental pathways and functional roles remain poorly understood. Using tissue mass cytometry, we localized B cells in the thymus and their orientation towards other cells in the medulla. We characterized the heterogeneity of thymic B cells using single-cell RNA sequencing of cells isolated from human thymi, identifying naive, germinal center-like, plasma cell, and multiple memory B cell populations. We identified a distinct pre-B cell subset with local thymic B cell development potential, that can develop due to the inhibition of Notch signaling by Deltex1, a Notch antagonist. Using BCR repertoire analysis, we explored clonal diversity, somatic hypermutation patterns and class switch recombination of thymic B cells. Spectral flow cytometry further validated the surface phenotype of thymic B cell populations and confirmed the presence of distinct CD21^-^CD27^-^ memory compartments with heterogeneous surface immunoglobulin isotype usage. In doing so we provide novel insights into the unique biology of human thymic B cells, their development, and their differentiation in the human thymus. We conclude that the human thymus supports local B cell development and differentiation into medullary memory-like populations that undergo class switching with limited somatic hypermutation, suggesting secondary lymphoid-like B cell programs adapted to central tolerance induction.

**One Sentence Summary:** The human thymus is not only a primary lymphoid organ for T cell development, but also a secondary site for the development and maturation of unique populations of B cells.

## INTRODUCTION

In the thymus, crosstalk between thymic stromal cells and developing thymocytes is required for the establishment of a diverse but tolerant T cell repertoire. To avoid autoimmunity, self-reactive T cells are deleted resulting in self-tolerant T cells. After positive selection in the thymic cortex, in which thymocytes with a functional T cell receptor (TCR) survive, thymocytes migrate from the cortex to the thymic medulla. Here, central tolerance is established through negative selection mediated by interactions with antigen-presenting cells (APCs) such as thymic dendritic cells and medullary thymic epithelial cells (mTECs). By presenting self-antigens, APCs ensure that thymocytes with autoreactive T cell receptors will be eliminated by clonal deletion or diverge to the regulatory T cell (Treg) lineage. (*1-3*).

Thymic B cells are a subset of APCs (*4-7*), making up a relatively small proportion of total thymic cells (∼0.5%) but already present in fetal life and increasing in numbers with age (*8*). While B cells are known to exist in the thymus (*9*) and to be potentially autoreactive (10), they have been less studied for their role as APCs compared to thymic dendritic cells and thymic epithelial cells and therefore less is known about their development and function in the thymus (*6, 9*). Thymic B cells are predominantly located in the medulla and express costimulatory molecules such as CD80, CD86 and high levels of MHC class II molecules establishing their role in negative selection (*4, 6, 7, 11*). To become activated, thymic B cells undergo a process called thymic B cell licensing by interaction with self-reactive SP CD4 cells through CD40L-CD40 signaling (*7*).

Licensing not only increases co-stimulatory capacity but can also induce a broader tolerogenic program in thymic B cells, including expression of the autoimmune regulator AIRE, linked to self-antigen display programs that are classically associated with medullary tolerance. This interaction between T and B cells will result in the isotype class switching of B cells intrathymically, enhancing their antigen presentation capabilities enabling negative selection (*7,12*). In addition to CD40L–CD40 signaling, type III interferon (IFN-λ) has been implicated in promoting thymic B-cell activation programs that support B cell–mediated induction of regulatory T cells (Tregs), further maintaining central tolerance (*7,13*). The importance of thymic B cells in the induction of Tregs has been demonstrated in mouse models deficient in thymic B cells, resulting in reduced numbers of thymic Tregs (*7, 13-15*). Recent studies have demonstrated that thymic B cells are essential for T cell tolerance induction to Aquaporin 4 (AQP4), a key autoantigen in the autoimmune disease neuromyelitis optica, confirming the importance of thymic B cells to the contribution of central tolerance (*16*). Lastly, the presence of class-switched antibody-producing thymic B cells was reported to be involved in mechanisms of central tolerance (*12*). Intrathymically differentiated CD138+ plasma cells that secrete immunoglobulins similar to natural antibodies (*17*) were described in human neonates (*18*), which may induce T cell tolerance towards dietary-specific antibodies.

We have previously shown that CD34^+^CDla^−^ thymocytes contain progenitors that preserve multi-lineage potential and found that the human thymus serves as the site where not only αβ and γδ T cells develop, but also non-T cells including NK cells, classical dendritic cells, plasmacytoid dendritic cells, monocytes, and B cells (*19, 20*). To better profile thymic non-T cells including the thymic B cells, we performed scRNA-seq on the combined subset of human thymocytes sorted for the expression of CD16, CD56, CD13, CD33, and CD19 that we here refer to as alternative lineage subset (i.e., non T cell lineage), and that we originally excluded from our analyses on T cell development subsets.

While the presence of B cells in the thymus and its role in central tolerance induction has been indicated, little is known about the differentiation of various B cell populations in the human thymus. In this study, we focus on examining B cells in the human thymus using scRNA sequencing, imaging mass cytometry and spectral flow cytometry. We investigated the heterogeneity of human thymic B cells and characterized multiple naïve, Germinal Center (GC)-like, plasma and memory B cell subsets including multiple subsets of CD21^−^atypical-like memory B cells. Using reconstructed BCR rearrangements we investigated isotype switching, clonal diversity and clonal overlap to gain more insight into intrathymic development, differentiation and maturation of B cells in the human thymus.

## RESULTS

### Characterization of non-T cell types in the thymus

We performed scRNA-seq on sorted thymocytes obtained from three healthy donors isolating the combined population of cells with expression of CD16, CD56, CD13, CD33, and CD19, referred to as the alternative lineage human thymocyte subset **(Fig. 1A)**. From this we generated scRNA-seq libraries for 5’ gene expression profiling using the lOx Genomics platform. In total, we obtained a dataset for approximately 10,000 cells obtained from three donors **(Fig. 1A; Suppl. Fig. 2A)**. From these gene expression libraries, we were also able to reconstruct BCR rearrangement data for almost every cell (see Methods).

**Figure 1.**
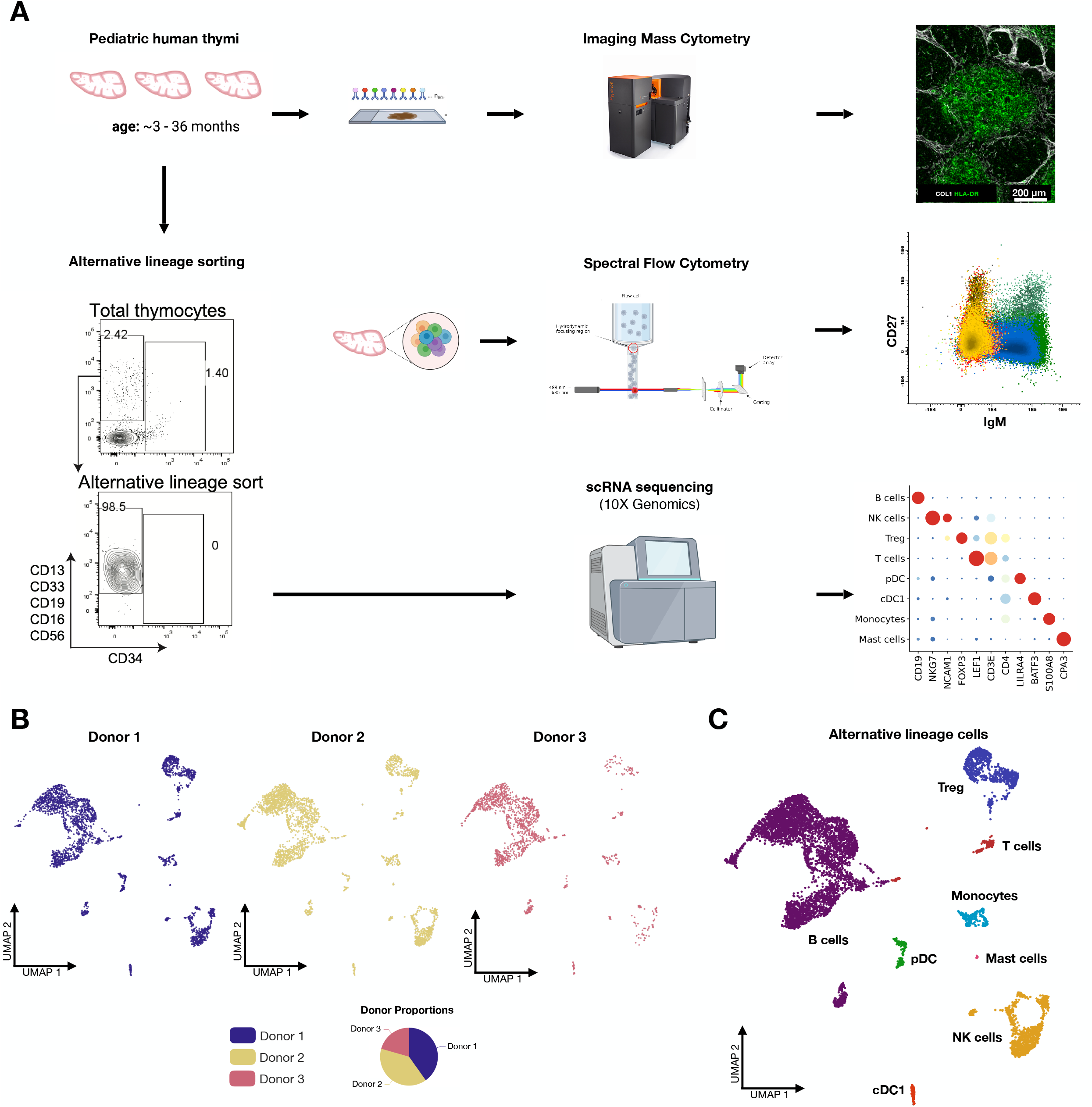
Multimodal profiling of human thymic alternative lineage cells reveals multiple non-T cell subsets. **(A)** Experimental scheme: Pediatric human thymic tissue was used for parallel multimodal analyses. Thymocytes from three donors were isolated, pooled and sorted based on expression of non-T lineage markers (CD13, CD33, CD19, CD16, and CD56). Sorted populations were sequenced by single-cell RNA sequencing for gene expression profiling. In parallel, total thymocytes from two donors were isolated and analyzed by spectral flow cytometry. Additional thymic tissue was analyzed by imaging mass cytometry. **(B)** UMAP projections showing the contribution of each of the three donors to the integrated single-cell dataset, with pie chart summarizing donor proportions. **(C)** UMAP of integrated alternative lineage cells annotated by cell identity, revealing multiple non-T cell populations including B cells, NK cells, cDCl, pDC, monocytes, mast cells, T cells, and Treg cells.

After *in silico* quality control to remove low-quality cells and doublets (*21*), we visualized the overall scRNA-seq data using a two-dimensional UMAP (uniform manifold approximation and projection) **(Fig. 1B)**. By manual annotation of our non-T cell dataset, we identified monocytes, cDCs, pDCs, mast cells, NK cells and B cells **(Fig. 1C)**.

### Thymic B cells mainly localize to the medulla in close interaction with SP thymocytes

While the population of B cells are the largest non-T cell type in our scRNA dataset **(Figure 1)**, unbiased spectral flow analysis showed that ∼1% of the total thymocyte population in the thymus constitute CD19^+^CD20^+^ B cells **(Fig. 1A; Suppl. Fig. 1)**. Tissue mass cytometry using the Hyperion Imaging system allows for the quantitative measurement of multiple (>35) antigenic markers using metal isotopes, enabling the identification of tissue heterogeneity with high precision. We used human thymic slices to investigate B cells **(Fig. 2C-F)**, plasma cells, and their localization in the context of developing T cells. As suggested by Spencer et al (*22*), B cells were primarily found in the medulla where they can make up to 30% of medullary cells in our analyses **(Fig. 2D)**. Our images showed that their frequency was indeed surprisingly high, approaching that of SP thymocytes **(Fig. 2D,F)**. Plasma cells were located primarily in the perivascular spaces between the cortex and medulla, as well as in the cortico-medullary junctions **(Fig. 2E)**, consistent with previous reports (*11, 18*).

**Figure 2.**
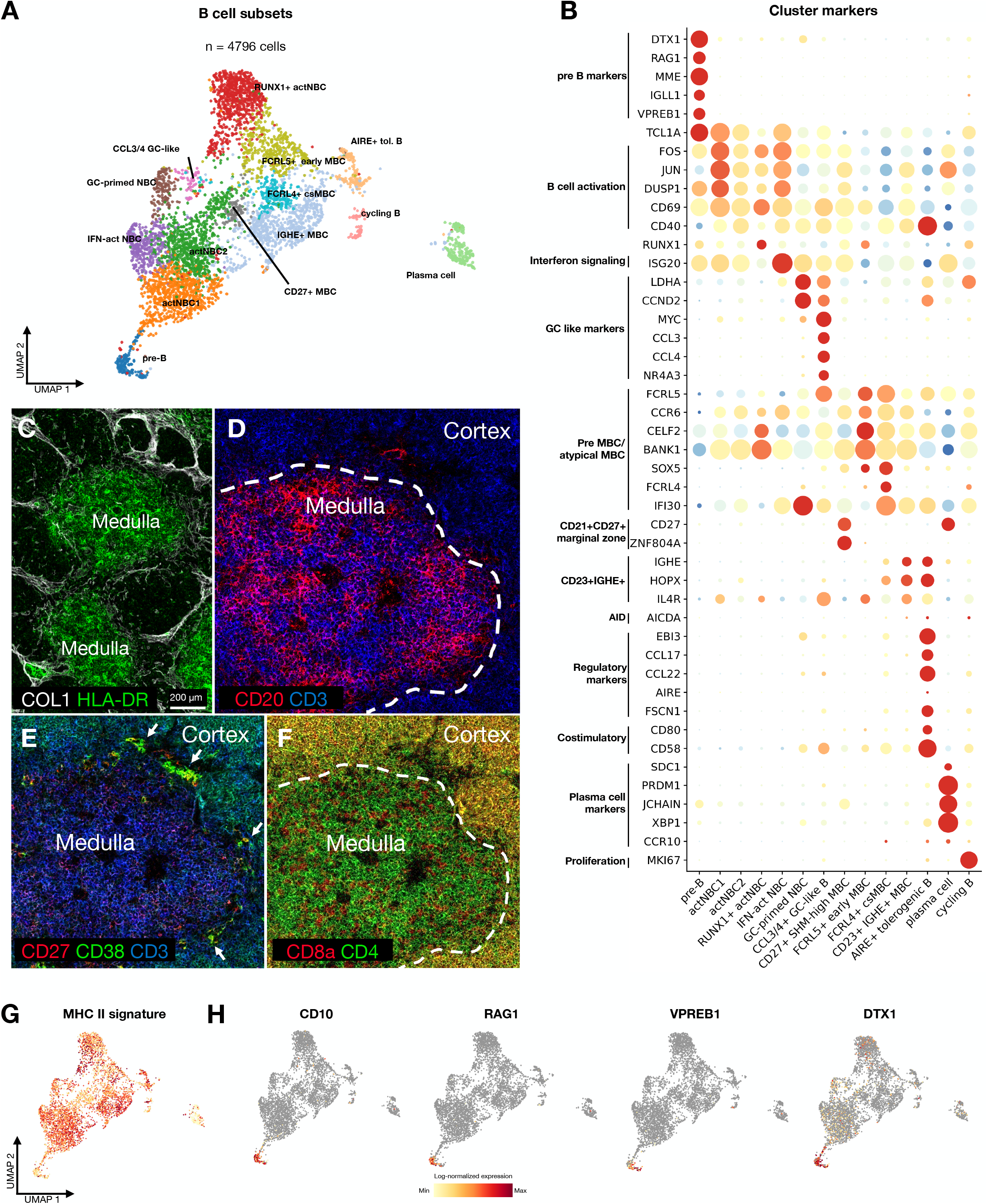
Single-cell RNA sequencing and histological analysis of the medulla in human thymi using tissue mass cytometry reveal heterogeneous B-cell populations which localize predominantly to the medulla. **(A)**UMAP projection of 14 transcriptomically distinct thymic B cell clusters. **(B)** Dot plot depicting the genes that identify thymic B cell clusters based on differential gene expression. **(C)** General overview of thymus medulla - cortex, HLA-DR showing highest concentration in the medulla. **(D)** Expression of B-cell marker CD20 (red) and T cell marker CD3 (blue) in the medulla and cortex. B cells are mostly located in the medulla. Dotted line, cortico-medullary junction. **(E)** Expression of Plasma cell markers CD27 (red), CD38 (green), and T cell marker CD3 (blue). Arrows depict plasma cells with expression of both CD27 and CD38. Plasma cells locate in the perivascular spaces and the cortico-medullary junctions. **(F)** Expression of CD8a (red) and CD4 (green) in the medulla and cortex. Single positive T cells localize in the medulla. Dotted line, cortico-medullary junction. **(G)** Combined expression (red is high combined expression) of MHC class II involved genes: *CIITA* (MHCII transactivator), *HLA-DRA, HLA-DRB1* (MHCII alpha and beta chain). **(H)** Projected expression of Notchl antagonist Deltexl (*DTX1*) and key markers of pre-B cells (red is high expression).

### Thymic B cells are a heterogeneous population and are different from classic subsets

To better investigate the heterogeneity of the thymic B cells we reanalyzed the B cell population containing around 5000 cells and generated a new UMAP plot. Clustering analysis revealed 14 sub-populations **(Fig. 2A)** which were annotated **(Fig. 2A,B)** using multiple scRNA tonsil (*23, 24*), and thymic B cell datasets (*23-26*). In addition to plasma cells, we identified naive B cells (NBC), multiple GC-like cell types and multiple memory B cell (MBC) populations, and a subset of pre-B (MME^+^IGHM^−^CD34^−^) cells **(Fig. 2B,H)**. Except for the pre-B cell and plasma cell populations, thymic B-cell subsets expressed CD40, consistent with their capacity to receive CD40L-dependent signals from interacting CD4 single-positive thymocytes **(Fig. 2B)**. While CD40 expression is a general feature of mature B cells, thymic B cells have been reported to display an unusually activated APC phenotype, characterized by high MHC class II and co-stimulatory molecule expression following intrathymic licensing. In line with this, thymic B-cell subsets showed high expression of MHC class II–associated genes: *CIITA* (MHCII transactivator), *HLA-DRA* and *HLA-DRB1* **(Fig. 2G)**, consistent with an APC-primed state described for licensed thymic B cells (*4, 7,12-14*).

### Thymic pre-B cells express Deltex1 that can negate the T cell fate by antagonizing Notch signaling

The thymic pre-B cell population shows clear differential expression of *MME* (CD10), *IGLL1, VPREB1* and *RAG1*. This population is significantly different from the rest of the thymic B subsets in the thymus (*27-31*) and resembles pre-B cells usually found in the bone marrow (32, 33) **(Fig. 2H)**. Notch signaling in the thymus would ordinarily repress the B cell fate of multi-potent progenitors (*34-37*) and push cells toward the T cell fate (38-40). In order to allow B cells to develop in the thymus Notch signaling has to be repressed, as shown by targeted mutation of *Notch1* in vivo resulting in abundant B cells intrathymically (*35, 36, 41). We* found that the thymic pre-B cells, just like pre-B cells in bone marrow, show high gene expression of *Deltex (DTX1*), a Notch target gene (*39, 42, 43*) that antagonizes Notch signaling (*44, 45*) in a negative feedback, providing an opportunity for B cell development from precursors that differentiate locally within the thymus environment (*46*) **(Fig. 2H)**. DN1 thymocytes also express *DTX1* in specific clusters coinciding with the upregulation of CD7 (*19*), suggesting thymic B cell development from thymus-seeding progenitors (*32, 47*) **(Suppl. Fig. 2B)**.

### Naïve B cells in the thymus are activated, licensed and engage early GC-associated programs

We identified multiple naïve B cell subsets (NBC) that together with the pre-B cell population are marked by the expression of *TCL1A*, which distinguishes these subsets from other B cells **(Fig. 2A,B)** (*48*). These clusters (actNBC1, actNBC2 and RUNX1^+^ actNBC) also have high expression of activation markers (*FOS, JUN, DUSP1 and CD69) (49*). In addition, we identified an interferon-activated naïve B-cell cluster (NBC IFN-act) enriched for interferon-stimulated genes such as *ISG20*, which has been linked to licensed B cells (13) **(Fig. 2B)**. In comparison to the other NBC clusters, NBC (3) shows a higher expression of RUNX1, which has been associated with resting B cells in mice (*50*). The fifth NBC cluster was characterized by high expression of *LDHA* **(Fig. 2B)**, a metabolic enzyme involved in glycolysis which has a specific role in the transition from naïve to activated B cells. Cells where LDHA was deleted failed to mount GC-dependent antibody responses (*51*). Also, Cyclin D2 (*CCND2*), a downstream target of MYC, is differentially expressed in this cluster. In secondary lymphoid organs, *MYC* is a well-established marker of strong help signals and selection in GC B cells, acting as a gatekeeper for further growth and expansion of a germinal center (*52-54*). Together this implies the fifth NBC as GC-committed. MYC itself is highly expressed in a cluster with differential expression of the proinflammatory chemokines *CCL3* and *CCL4* **(Fig. 2B; Suppl. Fig. 2C,D)**. These chemokines have been described to be secreted by activated and GC B cells, specifically elevated in centrocytes, B cells found in the light zone of a germinal center (*54, 55*). While centrocytes are described to interact with follicular dendritic cells and T follicular helper cells, Benet *et al* describe that secretion of CCL3/4 by B cells will attract Treg subsets responsive to these chemokines. Interestingly these cells also express *NR4A1-3* **(Fig. 2B)**, *NR4A1* is involved in differentiation of regulatory T cells (*56*) but in B cells is rapidly induced upon BCR signaling activation (*57*) and loss of naïve cell state (*58*). These cells together with the previously described cluster of GC-committed cells resemble an early GC-committed subpopulation also described as such in a scRNA atlas of tonsil cells by Massoni-Badosa *et al*. (*24*) **(Fig. 2B)**.

To validate the composition of the thymic B-cell compartment at the protein level, we performed spectral flow cytometry with a 19-color dedicated B cell panel. This showed that the majority of thymic B cells has a naïve phenotype (lgM^+^lgD^+^CD27^−^) and comprises approximately 82% of the total B-cell pool, whereas phenotypically defined memory B cells account for the remaining ∼18% **(Fig. 3A)**. Memory B cells were defined as CD27^+^ and/or class-switched based on immunoglobulin isotype staining **(Suppl. Fig. 1)**. Previous studies reported a prominent CD21^−^/low B-cell population in human thymus (*59*). Consistent with this, by investigating the expression of *CR2* (encoding for CD21), our scRNA-seq data suggested that thymic B-cell populations can be broadly separated intoCD21^+^ nai’ve/GC-like clusters and CD21^−^memory/plasma clusters **(Fig. 3A; Suppl. Fig. 3A)**, with the exception of a small CD21^+^CD27^+^ memory-like population **(Fig. 2A**,**3B)**. Spectral flow cytometry further refined this observation by showing that within the naïve B-cell compartment, ∼60%of cells were CD21^+^ and ∼40% were CD21^−^**(Fig. 3C)**, indicating that CD21 expression in thymic naïve B cells is not binary. Notably, CD21^−^/low naïve-like B cells displayed reduced surface IgM compared with CD21^+^ cells **(Fig. 3D)**, consistent with the idea that loss of CD21 accompanies activation of B cells within the thymic environment (*18*).

**Figure 3.**
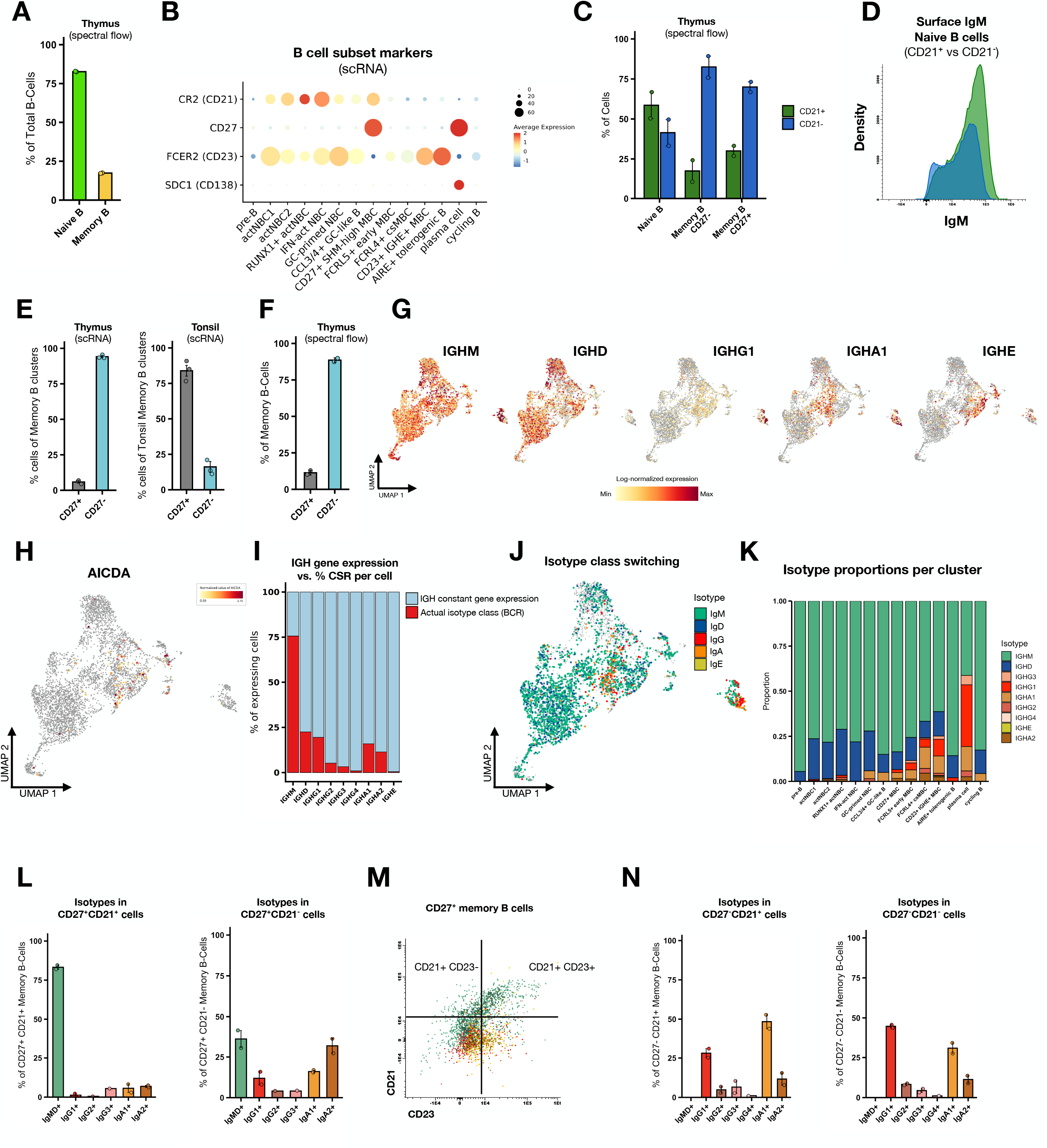
Most thymic memory B cells are CD21-low/negative, whereas the rare CD27^+^ memory compartment is largely unswitched. **(A)** Flow cytometry quantification of the thymic naïve (lgM^+^lgD^+^CD27^−^) and memory (CD27^+^ and/or IgA^+^/IgG^+^) B-cell compartments. **(B)** Dot plot showing expression of selected B-cell subset markers across thymic B-cell clusters, including CR2 (CD21), CD27, FCER2 (CD23), and SDC1 (CD138). (C) Flow cytometry quantification of CD21 surface expression within naïve, CD27^+^ and CD27^−^memory B-cell compartments. **(D)** Representative density plot of surface IgM expression in naïve thymic B cells comparing CD21^+^ (green) and CD21^−^(blue) populations. (E) Comparison of the frequency of CD27^+^ and CD27^−^cells within the memory B-cell compartment based on scRNA-seq data from thymus and tonsil. (F) Flow cytometry quantification of CD27^+^ and CD27^−^cells within thymic memory B cells. (G) UMAP feature plots showing expression of immunoglobulin heavy-chain constant region genes IGHM, IGHD, IGHG1, IGHA1, and IGHE across thymic B-cell clusters. **(H)** UMAP feature plot of AICDA gene expression. (I) Comparison between the proportion of cells expressing immunoglobulin heavy-chain constant region transcripts and the proportion of cells assigned as actually class-switched based on reconstructed BCR sequences. **(J)** UMAP projection colored by BCR-assigned isotype class. **(K)** Proportional distribution of BCR-assigned isotypes across thymic B-cell clusters. (L) Surface isotype composition of CD27^+^CD21^+^ and CD27^+^CD21^−^memory B cells measured by spectral flow cytometry. (M) Representative spectral flow plot of CD21 and CD23 expression within the CD27^+^ memory B-cell compartment, highlighting the presence of a CD21^+^CD23^−^subset. (N) Surface isotype composition of CD27^−^CD21^+^ and CD27^−^CD21^−^memory B cells measured by spectral flow cytometry.

### Thymic MBCs contain atypical-like MBC subsets with an antigen presentation gene signature

Having established that memory B cells represent a significant fraction of the thymic B-cell compartment, we next examined their heterogeneity in more detail. Spectral flow cytometry showed that within the phenotypically defined memory compartment, CD21^−^cells predominated, consistent with the enrichment of CD21^−^memory-like states in our scRNA-seq dataset **(Fig. 3C)**. In particular, within class-switched CD27^−^memory B cells, approximately80% of cells lacked CD21 surface expression, whereas only ∼20% retained CD21 **(Fig. 3C)**. These data indicate that the dominant CD27^−^memory compartment in thymus is largely CD21^−^. Within our scRNA-seq data we identified 4 different subsets of CD21^−^CD27^−^cells, which resemble the atypical MBCs described by King *et al* in human tonsils **(Fig. 3E)**. Atypical CD20^+^CD21^−^/lo/CD27^−^memory B cells have been described as more tissue resident-like cells (*60-62*). Previously these CD21^−^/lo cells have been described in the infant thymus as APCs (59) with potential for increased T cell-B cell interactions (*63*), which fits with the upregulation of genes involved in antigen presentation described by King et al. Multiple studies have shown that CD21^−^tissue-like memory or atypical memory B cells can be defined by the expression of Fc receptor-like (FCRL) 4 and 5 (*60-62*), which are negative regulators of B cell receptor signaling (*60, 64*). The role of these atypical memory B cells remains elusive, but some studies link these subsets to autoimmunity (*65, 66*). Indeed, cells without CD21 expression have high expression of *FCRL5* in our dataset **(Fig. 2B, Suppl. Fig. 2F)**. Of the naïve B cell clusters, RUNX1+ actNBC and GC-primed NBC cells show some expression of *FCRL5*, while still being positive for CD21 (*CR2*) **(Fig. 2A,B; Suppl. Fig. 2E)**. The RUNX1+ actNBC cluster also resembles a pre-MBC population described by Holmes *et al*. marked by the expression of *BACH2, CCR6, CELF2* and *BANK1* in their tonsil data (*26, 67*) **(Fig. 2A,B)**. RUNX1+ actNBC seems to be transcriptionally connected to a cluster we labeled FCRL5+ early MBC **(Fig. 2A,B)**. These cells also have high expression of *FCRL5* combined with the expression of the transcription factor SOX5 **(Fig. 2B; Suppl. Fig. 2F)**, which is characteristic of specific stages of B cell differentiation of activated B cells and in atypical memory cells with low CD21 expression (*61, 64, 68*). Furthermore, these cells start to induce IgA transcripts as shown by increased expression of *IGHA* **(Fig. 3G)**, leading to IgA class switching **(Fig. 3J,K)**, most likely by CD40 engagement (*69, 70*). We also classified a cluster that differentially expressed *FCRL4*, which we classified as class-switched MBC (FCRL4+ csMBC) **(Fig. 2A,B; Suppl. Fig. 2F)**. Cells in this cluster continue to have high expression of *SOX5* and *IGHA1* and showed IgG class switching of their BCR **(Fig. 3G,J,K)**. This cluster fits the description of Ehrhardt *et al*. that *FCRL4* expression is restricted to a subpopulation of memory B cells with increased expression of the costimulatory molecules CD80 and CD86 suggesting a role as antigen-presenting cells (*11*) **(Fig. 2B; Suppl. Fig. 2E)**. Additionally, these cells have high expression of GILT (*IFI30*), which is also highly expressed in APCs (*71*) **(Fig. 2B; Suppl. Fig. 2E)**.

### Thymic CD27+ memory B cells are rare and mostly unswitched (lgM^+^)

Whereas memory B cells in secondary lymphoid organs are typically enriched for a CD27^+^ compartment (*62*), thymic memory B cells in our scRNA-seq dataset showed limited CD27 expression: only ∼10% of transcriptomically defined memory B cells expressed CD27 mRNA **(Fig. 3E)**. To assess whether CD27 transcript detection reflects CD27 surface expression in the thymic setting, we quantified CD27 protein by spectral flow cytometry **(Fig. 3F)**. Memory B cells were defined phenotypically as CD27^+^ and/or class-switched based on immunoglobulin isotype staining **(Suppl. Fig. 1)**. Within this memory gate, ∼10% of cells expressed CD27 at the cell surface **(Fig. 3F)**, closely matching the fraction of CD27^+^ cells observed at the mRNA level in scRNA-seq **(Fig. 3E)**. While mRNA-protein concordance varies by marker, these data suggest that for CD27 in this context, transcript detection provides a good estimate of cell surface expression. This low frequency of CD27^+^ memory B cells in thymus contrasts with published scRNA-seq data from human tonsil, where CD27 gene expression is seen in the majority of memory B cells (approximately ∼80%) (Massoni-Badosa, Aguilar-Fernândez et al. 2024) **(Fig. 3E; Suppl. Fig. 3D)**, highlighting an important phenotypic difference of the thymic memory-like B-cell compartment compared to secondary lymphoid organs.

Within the scRNA-seq data, the isotype features of these few CD27^+^ thymic B cells were consistent with a predominantly unswitched identity. At the transcript level, cells expressed IGHM and IGHD, with occasional detection of class-switched constant region transcripts **(Fig. 3G,H,I)**. To assess isotype usage more directly, we used reconstructed BCR sequences to assign heavy chain constant regions. This confirmed that the majority of BCRs in this cluster were IGHM and/or IGHD, with only a small fraction assigned to class-switched isotypes including IGHA1, IGHA2, and IGHG1 **(Fig. 3G,H,I)**. The spectral flow data of the previously mentioned thymic CD27+ population was used to validate this isotype composition. Within this memory compartment, the majority of cells were CD21^−^(∼70%), with a smaller CD21^+^ fraction (∼30%) **(Fig. 3C)**. These two compartments showed significant differences in surface isotype composition: CD27^+^CD21^+^memory B cells were predominantly non-class-switched (lgM/lgD^+^) **(Fig. 3L)**, whereas CD27^+^CD21^−^memory B cells contained substantially higher proportions of class-switched cells, including IgA- and IgG-expressing populations **(Fig. 3L)**. Together, these data indicate that unswitched CD27^+^ memory-like cells retain CD21, while class-switched CD27^+^ memory cells are enriched within the CD21^−^compartment **(Fig. 3L; Suppl. Fig. 1)**. Notably, this cluster displayed the highest somatic mutation frequency among thymic B-cell subsets **(Fig. 4)**, consistent with the increased mutation burden typically observed in CD27^+^ memory B cells in tonsil (King, Orban et al. 2021). Functionally, the presence of a highly mutated CD27^+^ unswitched memory-like subset in thymus is notable because in adults, mutated and selected CD27^+^ memory B cells can rapidly differentiate into antibody-secreting cells upon restimulation (72, 73).

**Figure 4.**
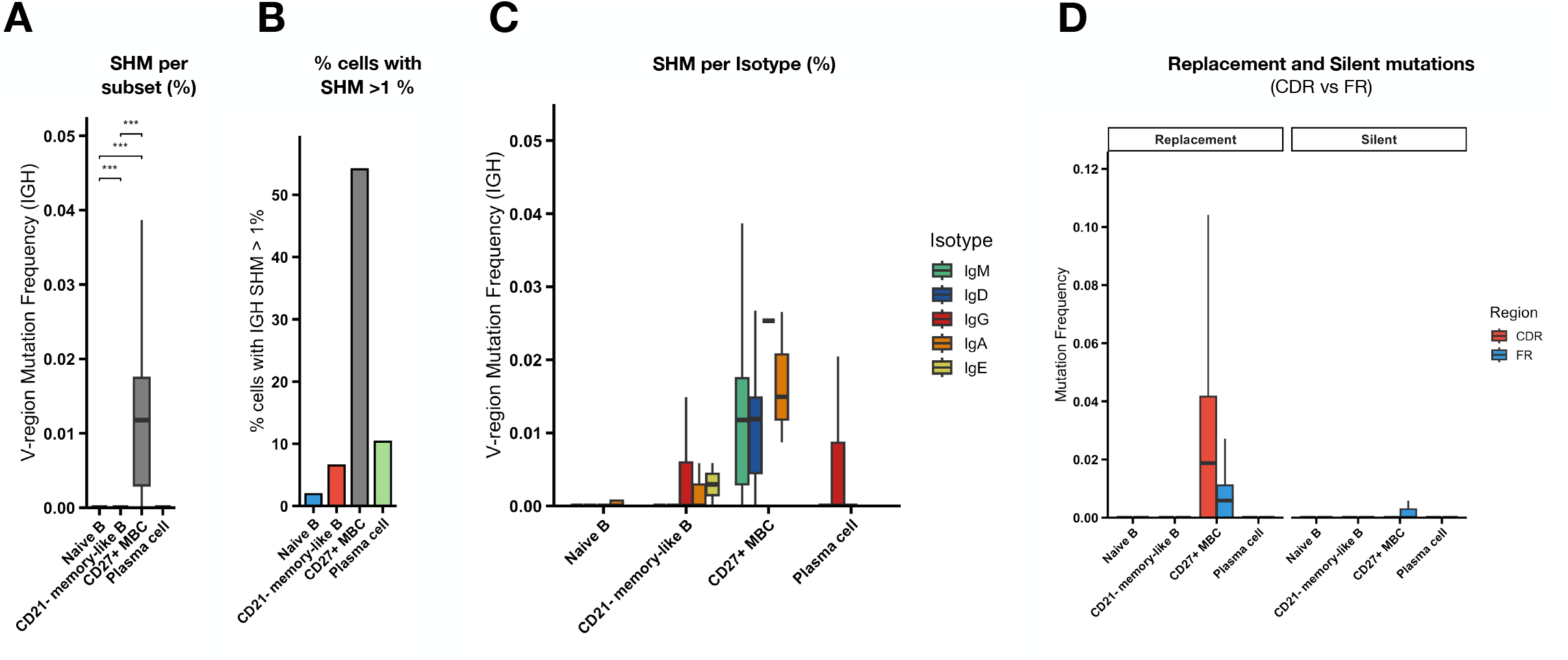
Low somatic hypermutation in class-switched thymic B cells suggests GC-independent differentiation. **(A)** Distribution of IGH V-region mutation frequencies across the four major thymic B-cell compartments: naïve B cells, CD21^−^memory-like B cells, CD27^+^ memory B cells, and plasma cells. SHM frequencies were calculated as the number of nucleotide mutations in the IGH V region divided by the total number of nucleotides analyzed. **(B)** Fraction of cells in each compartment with >1% IGH SHM, containing a substantial fraction of mutated cells. **(C)** IGH SHM frequencies displayed by BCR-assigned isotype within the four major thymic B-cell compartments. **(D)** Distribution of replacement and silent mutations in complementarity-determining regions (CDR) and framework regions (FR) across the four major thymic B-cell compartments, showing enrichment of replacement mutations in the CDR of CD27^+^ memory B cells, consistent with antigen-driven selection.

**The human thymus contains marginal zone-like CD21*CD23’CD27*lgM* memory B cells**

Perera *et al* described that in mice thymic B cells are composed of CD21^+^CD23^+^, resembling follicular B cells or CD21OD23’ B cell subsets, but that the mouse thymus does not contain CD21^+^CD23^−^marginal zone memory B cells (MBCs). We found that the CD27^+^ memory B cell cluster indeed was positive for the expression of CD21, while negative for CD23 (by investigating the expression of genes *CR2* (CD21) and *FCER2* (CD23)) **(Fig. 3B)**. We next examined CD23 expression within the CD27^+^CD21^+^ memory-like compartment by spectral flow cytometry. CD23 surface expression was heterogeneous and spanned a continuum, including a fraction of CD23^^^low/negative cells within the CD27^+^CD21^+^gate which are mostly lgM^+^ **(Fig. 3M)**. This suggests that, different from mouse, the human thymus contains an unswitched CD21^+^CD23^−^CD27^+^ subset resembling marginal zone-like B cells. lgM^+^lgD^+^CD27^+^ (IgM memory’) B cells are a well-described recirculating population in human blood linked to splenic marginal zone biology, shown to carry somatic mutations, and mutated antibodies (*74, 75*). Their presence in thymus is consistent with a contribution of circulating memory-like B cells to the thymic B-cell pool.

### Thymic B cells express AID and undergo heterogeneous class switching

Next to differential gene expression analysis, the thymic B subsets could be separated in class-switched and non-class-switched subsets based on the expression of the different genes encoding for Ig class, combined with the reconstructed BCR data **(Fig. 3G,J; Suppl. Fig. 3F;** see Methods). Isotype class associated IGH constant region genes *IGHM* (IgM) and *IGHD* (IgD) are expressed in all clusters (apart from *IGHD* in plasma cells) **(Fig. 3G)**. Expression of IGHG, IGHA and IGHE is differentially expressed across the memory B cell subsets **(Fig. 3G)**. High expression of *IGHG1* is limited to plasma cells while *IGHG2-4* are widely expressed across the MBC clusters **(Fig. 3G; Suppl. Fig. 3B)**. The expression of *IGHA1-2* and *IGHE* is mutually exclusive between the MBC clusters **(Fig. 3G; Suppl. Fig. 3B)**. These clusters express activation-induced cytidine deaminase (AID) encoded by the gene *AICDA* **(Fig. 3H)**, suggesting that class switch recombination (CSR) and somatic hypermutation (SHM) can occur in the human thymus. Expression levels of AICDA found in our thymic MBC clusters are similar to the levels observed in MBCs in human tonsil scRNA-seq data **(Suppl. Fig. 3E)**. In mice, AID expression in thymic B cells was shown to be ∼13−24% lower than in B cells isolated from spleen and lymph node (12), and AID expression can vary between anatomical sites and even between different regions of the germinal center (76, 77).

While the expression of IGH constant region genes is associated with an isotype class, it does not necessarily indicate actual class switching of a B cell (78). Cells can also produce germline transcripts coming from noncoding transcripts of the constant immunoglobulin genes, which could indicate priming for future class switching of isotypes (78). For definitive isotype class identification within our scRNA-seq data, we used the class identified from the reconstructed BCR data. This showed that the frequency of class-switched B cells is indeed lower compared to the expression of the IGH constant region genes **(Fig. 3I)**. As expected, naïve B cells have a low proportion of switched cells which starts to increase in early memory B cells with cells switching to IgG or IgA **(Fig. 3J,K)**. The highest number of switched cells are mature MBCs (∼25%) and especially in plasma cells (60%) where most cells have an IgG isotype **(Fig. 3K)**.

To validate isotype usage at the protein level, we quantified surface immunoglobulin isotypes by spectral flow cytometry **(Fig. 3L; Suppl. Fig. 1)**. Consistent with BCR-based constant region assignments, the majority of thymic B cells displayed a non-class-switched phenotype (IgM/IgD) **(Suppl. Fig. 3G)**, whereas class-switched cells (IgA and/or IgG) represented a smaller fraction of the total compartment **(Suppl. Fig. 3G; Suppl. Fig. 1)**. Among class-switched CD27^−^memory B cells, lgG1 and lgA1 are the dominant surface isotypes, with smaller contributions from lgA2 and lgG2/lgG3 and minimal IgG4 **(Fig. 3N)**. CD21+ and CD21-memory B cells show a shift in subclass composition: the CD27^−^CD21^−^ fraction was slightly enriched for lgGl, whereas the CD27^−^CD21^+^fraction was enriched for lgAl **(Fig. 3N)**. Representative biaxial plots illustrating the gating strategy for IgA and IgG subclasses are shown in **Suppl. Fig. 1**. Together, these protein-level data support the BCR-based constant-region assignments and indicate that class-switched thymic memory cells are heterogeneous in isotype composition, with CD21 status marking distinct compartments.

### Low somatic hypermutation in class-switched thymic B cells suggests GC-independent differentiation

To further examine diversity and affinity maturation of the thymic B cells we also used the BCR rearrangements to quantify IGH somatic hypermutation (SHM) frequencies **(Fig. 4; Suppl. Fig. 4)**. Across the thymic B-cell compartment the overall mutation frequency is low for all subsets (∼0% -1.2%) **(Fig. 4A)**. Because overall SHM values were highly zero-inflated across thymic B-cell subsets, we additionally quantified the proportion of cells with >1% IGH SHM to distinguish compartments containing a substantial fraction of clearly mutated cells from those in which the observed SHM signal was driven by only a few highly mutated cells **(Fig. 4B)**. This showed that SHM was strongly concentrated in the CD27^+^ memory B-cell compartment, in which both the overall mutation frequency and the proportion of cells with >1% IGH SHM were markedly higher than in the other groups **(Fig. 4A,B)**. In contrast, naïve B cells showed only rare, mutated cells, while CD21^−^memory-like B cells contained only a small fraction of mutated cells and remained close to germline overall **(Fig. 4A,B)**. Plasma cells likewise showed low overall SHM, with only a minority of cells exceeding the 1% threshold **(Fig. 4A,B)**. We also compared SHM by isotype within the major thymic B-cell compartments **(Fig. 4C)**. Again, the highest mutation frequencies were observed in the CD27^+^ memory compartment, particularly among unswitched IgM^+^ and IgD^+^ cells. In contrast, CD21^−^memory-like B cells and plasma cells contained class-switched IgG^+^/IgA^+^ cells with comparatively low SHM, indicating that in the thymus class switching can occur without the usual high rates of somatic hypermutations. To further examine whether the mutations in CD27^+^ MBCs showed evidence of antigen-driven selection, we compared replacement and silent mutation frequencies in CDR and framework regions **(Fig. 4D)**. Replacement mutations were enriched in the CDR relative to the FR regions, whereas silent mutations were rare, which is consistent with selective pressure on the antigen-binding region.

### Thymic B cells display clonal diversity and clonal overlap across subsets, suggesting these subsets arise from a common ancestral lineage

Next, we analyzed the B cell receptor (BCR) repertoires across our thymic B cell subsets to investigate clonal diversity and clonal relationships **(Fig. 5)**. All of the B cell subsets show a high proportion (>90%) of unique clones **(Fig. 5A)** suggesting a high clonal diversity. While the number of clonotypes varies because of the difference in size between subsets, the distribution and frequency of clonotypes across subsets does not indicate clonal expansion **(Fig. 5B)**. To further assess clonal diversity of our BCR repertoires we calculated Shannon indices across subsets **(Fig. 5C)**. The Shannon diversity index shows minimal differences between the individual indices of every subset, as well as minimal differences between naïve, CD21-memory-like, CD27+ MBC and plasma cells with indices ranging from 3.63-3.72 indicating an evenness of repertoires (*79-83*) **(Fig. 5C)**.

**Figure 5.**
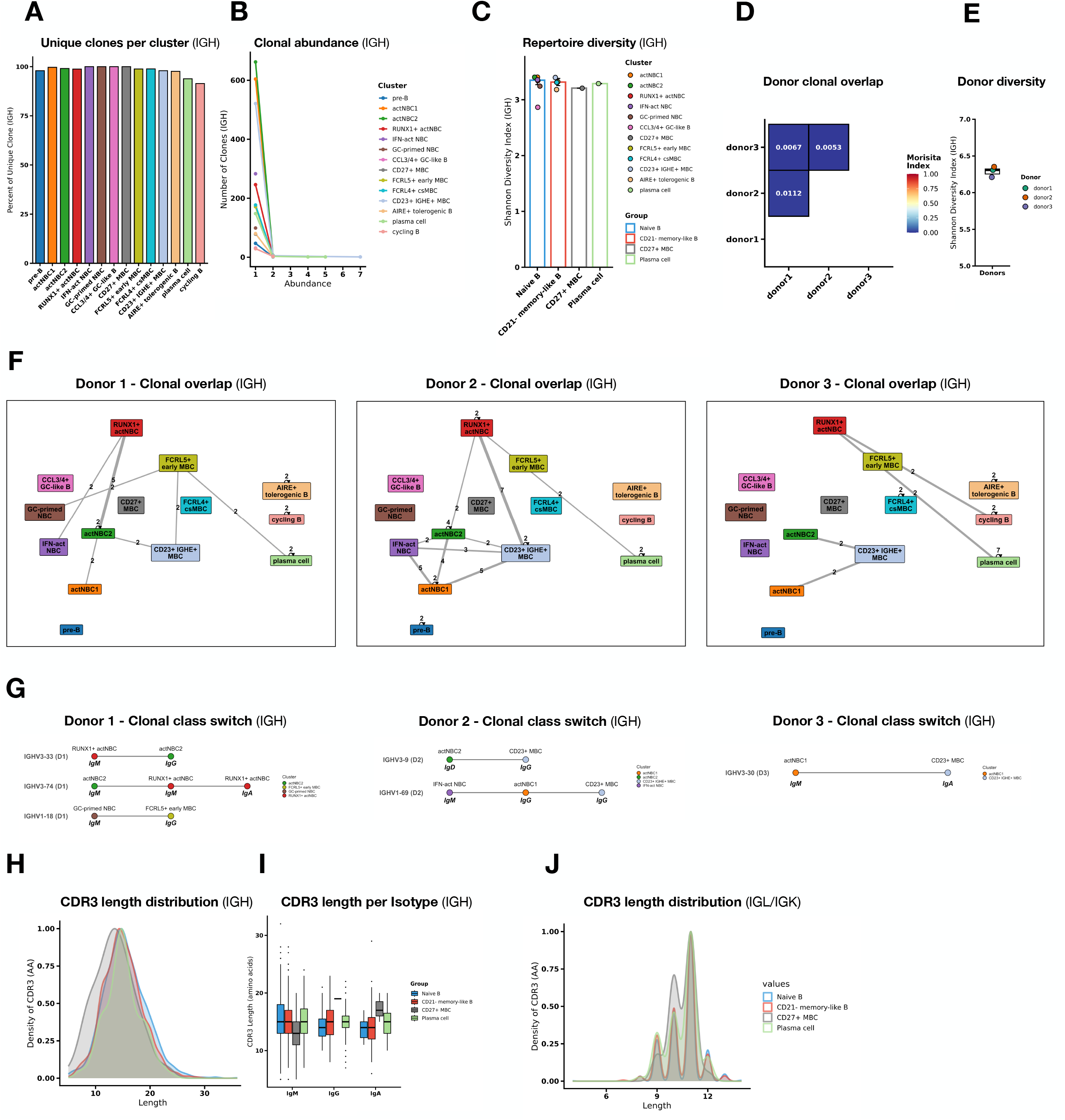
Thymic B cells display clonal diversity and clonal overlap across subsets, suggesting a lineage relationship. **(A)** Proportion of unique IGH clones within each annotated thymic B-cell cluster. **(B)** Relative abundance (number of occurrences of clonotypes) distribution of IGH-defined clonotypes across thymic B-cell clusters. **(C)** Repertoire diversity measured by Shannon diversity index of each annotated thymic B-cell cluster grouped by the four major thymic B-cell compartments (Naïve B, CD21^−^memory-like B, CD27^+^ MBC, and plasma cells). **(D)** Donor-to-donor clonal overlap based on IGH repertoires, calculated using the Morisita-Horn index, showing minimal overlap between donors. **(E)** Shannon diversity of the donor-specific IGH repertoires. **(F)** Within-donor IGH clonal overlap networks for each donor, showing shared clone groups between multiple thymic B-cell clusters. Edge thickness reflects the number of overlapping clones. **(G)** Within-donor IGH clonal class-switch networks showing clones overlapping with unswitched and class-switched states across transcriptomically different thymic B-cell clusters. **(H)** Density plot of IGH CDR3 amino-acid length distributions across the four major thymic B-cell compartments. **(I)** Boxplot depicting differences in IGH CDR3 amino-acid length for IgM, IgG and IgA between the four major thymic B-cell compartments. **(J)** Density plot of IGL CDR3 amino-acid length distributions across the four majorthymic B-cell compartments.

Additionally, we investigated clonal relationships within the thymic BCR repertoire. As a first step, we calculated the clonal overlap between donors using the Morisita-Horn index, based on IGH and IGL repertoires separately (*84*) **(Fig. 5D)**. Clones were defined by shared V(D)J gene usage and CDR3 amino-acid similarity using a 0.8 Levenshtein threshold (see Methods). Only minimal overlap was detected between donors, with Morisita-Horn indices of 0.005-0.01 for IGH, indicating that each donor harbored a largely private B-cell repertoire. This was consistent with the similar Shannon diversity indices observed across donors **(Fig. 5E)**, suggesting comparable overall repertoire diversity despite limited clonal sharing. Similar analyses based on IGL showed broader overlap both between donors and across clusters **(Suppl. Fig. 5A-E)**, consistent with the lower repertoire complexity of the light chain; therefore, subsequent interpretation of clonal continuity and class-switch relationships was based primarily on IGH.

Because clonotypes are donor-specific, all cluster-level overlap analyses were performed within individual donors to avoid dilution of biologically meaningful clonal relationships by unrelated repertoires. Although infrequent, shared clones were detected between multiple clusters representing distinct developmental states, supporting clonal relationships between activated naïve, memory-like, and plasma-cell subsets. Clonal overlap was observed between activated naïve B-cell clusters, FCRL5^+^ early memory B cells, and the CD23^+^ IGHE^+^ memory-like cluster, connecting a transcriptionally different but clonally connected differentiation trajectory **(Fig. 5F)**. More importantly, we identified clones connecting unswitched and class-switched states within individual donors. These included donor-specific clones linking IgD^+^ actNBC2 cells to IgG^+^ CD23^+^ memory-like cells, as well as clones connecting IgM^+^ IFN-act NBCs, lgG^+^ actNBCl cells, and IgG^+^ CD23^+^ memory-like cells **(Fig. 5G)**. The presence of shared IGH clones across phenotypically different unswitched and switched compartments supports lineage relation between these thymic B-cell subsets and is consistent with local class-switching during intrathymic differentiation. Of note, the CD27^+^ memory B-cell compartment did not show appreciable clonal overlap with the other thymic B-cell subsets, which further supports the idea that these cells represent a subset of B cells biologically distinct from the other thymic B cells subsets **(Fig. 5F)**, suggesting they may possibly have originated from outside the thymus.

Finally, we examined the distribution of complementarity-determining region 3 (CDR3) amino acid lengths of the reconstructed BCRs across the major thymic B-cell compartments. The CDR3 region forms the central part of the antigen-binding site of the immunoglobulin and is generated during V(D)J recombination by nucleotide addition and deletion at the junctions of the rearranging gene segments (85, 86). A normal distribution of CDR3 lengths reflects the expected diversity and adaptability of the immunoglobulin repertoire in health and disease (*87, 88*).

Analysis of CDR3 length distributions showed broadly normal and highly overlapping profiles across the four major thymic B-cell groups for both IGH and IGL/IGK chains. As expected, IGH CDR3s displayed a broader length distribution than light-chain CDR3s, whereas IGL/IGK CDR3s were tightly constrained across all groups **(Fig. 5H-J; Suppl. Fig. 5J,K)**. For IGH, naïve B cells, CD21^−^memory-like B cells, and plasma cells all showed very similar median CDR3 lengths of 15 amino acids, with mean lengths ranging from 15.0 to 15.5 amino acids. In contrast, the CD27^+^ memory B-cell compartment showed a modest shift toward shorter heavy-chain CDR3s, with a median length of 13 amino acids and a mean of 13.5 amino acids **(Fig. 5H)**. For IGL/IGK, all four groups were highly similar, with median CDR3 lengths of 9–10 amino acids and only minimal variation in mean length and spread **(Fig. 5J)**. When IGH CDR3 length was further stratified by BCR-assigned isotype, most isotype groups remained broadly overlapping across the four major thymic B-cell compartments **(Fig.51)**. The clearest difference was observed in the CD27^+^ MBC compartment, in which IgM-associated CDR3s were shorter overall than in the other groups. In contrast, IgG and IgA CDRS lengths did not show a consistent shift across the dominant thymic memory-like compartments, although some variation was present in the smaller isotype-specific subsets **(Fig. 51)**.

### Identification of CD23^+^lgA/lgG^+^ cells ready to switch to IgE

While thymic atypical MBCs do not express CD21 and CD23, as described, we also identified B cells with expression of interleukin-4 (IL-4)–and IL-13–regulated genes such as *IL4R, FCER2* (CD23) **(Fig. 2B, 3B)** and while the BCR data showed that these cells express IgA or IgG **(Fig. 3J,K)**, they show differential expression of *IGHE* **(Fig. 3G)**. At the transcriptome level, this cluster shows close resemblance to CD27^+^CD23^+^IgG1^+^ memory B cells, termed type 2 memory B cells, which were found enriched in children with peanut allergy from whom CD27^+^ memory B cells were sequenced using scRNA (89). This study described that type 2 memory B cells transcribing *IGHE* will most likely differentiate into IgE producing plasma cells. Without CD27, the here described CD23^+^lgA/lgG^+^ cells are phenotypically different from the subsets described by Ota *et al (89*). Transcriptomically on the other hand, there is a large overlap between the differentially expressed genes, with genes such as *IGHE, HOPX, IL4R* and *S100A10* that predominantly overlap with our data **(Fig. 2B)**. These genes together with genes involved in the MHC class II pathway such as *HLA-DPB1, HLA-DP1, HLA-DQA1* **(Suppl. Fig. 2E)**, suggest that these cells have increased antigen-presenting capabilities. The discrepancy between the expression of the genes of the immunoglobulin constant regions and the class detected on the BCRs was also something Ota *et al* noticed, which is especially striking when comparing the constant region gene expression of *IGHE* and the BCR-assigned heavy-chain isotype for IgE **(Suppl. Fig. 3H)**. The gene coding for *IGHE* is differentially expressed in multiple MBC-like clusters **(Fig. 2B, 3G)**, while only two cells were classified as IgE class-switched from the BCR data **(Fig. 3I,J,K; Suppl. Fig. 3H)**. Interestingly, compared to all called isotypes cells, in cells grouped by BCR-assigned isotype, B cells with isotype IgG are the only cells in which *IGHE* is highly expressed next to *IGHG1-4* **(Suppl. Fig. 3H)**.

### AIRE expressing thymic B cells resemble regulatory cell types

The second, much smaller, cluster enriched for *IGHE* is also positive for CD23 but differs from the previous cluster by the differential expression of *EBI3* and *CCL22, CCL17*, which can recruit CCR4-expressing cells to the medulla, such as Tregs (*90*) **(Fig. 2A,B; Fig. 3B; Suppl. Fig. 2D)**. Cells in this cluster also display increased expression of genes in the CD40 signaling pathway (including *CD40, TRAF1*, and *ICAM1*), costimulatory molecules like CD80 and CD58 (*91*) and NF-κB activation (including *NFKBIA, NFKB1, NFKB2*) **(Suppl. Fig. 2G)**, most likely by T cell-mediated activation and antigen stimulation (*26*).

Both Cordero *et al* and Heimli *et al*. described a similar small cluster of human thymic B cells within scRNA-seq data within CD21^−^CD35^−^thymic B cells (18) and APC-enriched cells (25). Heimli *et al*. described resemblance to aDC2, while Cordero *et al* found comparisons with regulatory/tolerogenic cells based on the expression of *EBI3 (92). We* also found overlap with multiple mregDC markers like *EBI3, CCL17, CCL22* and *FSCN1* **(Fig. 2B)** (93), confirming possible regulatory capabilities of these cells. This cluster was notably the only cluster with AIRE expressing cells **(Fig. 2B; Suppl. Fig. 2H)**. AIRE expression by thymic B cells has been described as a thymus-specific feature and has been linked to presentation of AIRE-dependent self-antigens, supporting clonal deletion of autoreactive CD4 single-positive thymocytes and central tolerance induction (*7, 33*). Together, these features suggest that this rare thymic B-cell subset represents an activated, tolerogenic APC-like population positioned to promote thymocyte tolerance through recruitment/interaction cues and AIREassociated self-antigen presentation.

### Thymic plasma cells express gut specific chemokine receptor CCR10 and multiple isotype classes

Plasma cells are a distinct cluster within the UMAP **(Fig. 2A)**, these CD21^−^CD23^−^CD27^+^ B cells have high expression of plasma cell markers CD138 (*SDC1*), BLIMP-1 (*PRDM1*),*JCHAIN*, and *XBP1*, which closely resembles scRNA data from previously published human thymic plasma cells (18) **(Fig. 2B)**. Consistent with this identity, Imaging mass cytometry (Hyperion) identified a CD27^^^hiCD38^^^hiCD20^^^low/-population consistent with antibody-secreting cells that localized predominantly to COL1^+^ perivascular spaces (PVS) **(Fig. 2E)**, providing spatial validation of the thymic plasma cell compartment. These cells have low expression of genes involved with MHC class II compared to the other clusters **(Fig. 2B, Suppl. Fig. 2E)** and overexpression of genes related to immunoglobulins indicating they are dedicated to antibody production **(Fig. 3G, Suppl. Fig. 3B)**. BCR constant-region data indicated that most thymic plasma cells were class-switched, with IgG as the predominant isotype followed by IgA, while a smaller fraction that retained IgM/IgD constant regions **(Fig. 3J,K)**. This was reflected by expression of *IGHG1-4, IGHA1 -2*, and *IGHM* transcripts **(Fig. 3G, Suppl. Fig. 3B)**. Although IGHE transcripts were detectable in a subset of plasma cells, IgE constant-region calls were rare, consistent with the broader observation that IGHE transcription can occur without IgE switching **(Fig. 3G,J,K; Suppl. Fig. 3H)**. Furthermore, there are some cells from the CD23^+^IGHE^+^ cluster that overlap with the plasma cells **(Fig. 2A)**, suggesting these cells could be precursors to thymic plasma cells. Thymic antibody-secreting cells including plasma cell populations have been previously described in mice and humans (*11, 66, 94, 95*) including their possible intrathymic development (*18*) and their involvement in dietary tolerance (*17, 96*). In addition, some of the plasma cells expressed CCR10 **(Fig. 2B, Suppl. Fig. 2C)**, a chemokine receptor also described being upregulated on IgA^+^ plasma cells generated in the gut lymphoid tissue in order to migrate into the colon for colonic homeostatic regulation (*97*). This suggests that a fraction of thymic antibody-secreting cells may acquire tissue-homing programs typically linked to mucosal immunity.

## DISCUSSION

The thymus is traditionally viewed as a primary lymphoid organ, responsible for the development and maturation of T cells. During T cell development T cells acquire tolerance to self-peptides and during this process autoreactive clones are eliminated by clonal deletion or redirected to develop into regulatory T cells. While often overshadowed by DCs, B cells comprise a substantial proportion of the thymic APC pool. Our data show that the thymic B cell compartment is not a collection of imported cells from the periphery but a dynamic, differentiating lineage that plays a unique role in tolerance induction (*7*) including cells that develop intrathymically as well as mature B cells that can enter the medulla region. Using a multimodal approach, our study sheds new light on the functional and transcriptional diversity of human thymic B cells. With mass cytometry we localized thymic B cells and with scRNA-seq we characterized distinct thymic B cell subsets, including naïve, germinal center (GC)-like, memory and plasma cells, along with atypical-like memory B cell subsets and regulatory B cells. Combined with BCR profiling and spectral flow analysis we demonstrate that thymic B cells are a unique subset of B cells which can develop and which display differentiation capacity from naïve to memory B subsets with an antigen-presenting phenotype.

One pathway of origin could be that thymic B cells develop from fetal liver and bone marrow pre-B cells that migrate to the thymus, or that B cells differentiate from uncommitted thymic seeding progenitor (TSP) cells (*32, 47, 98*). In both cases, Notch signaling must be blocked in the thymus to prevent cells from being directed to the T cell fate. Deltex1 is an E3 ubiquitin ligase that interacts with the intracellular domain of the Notch receptor and in this manner can inhibit Notch signaling by directing the Notch receptor towards lysosomal degradation thus preventing downstream signaling (*44, 45*). Fetal thymic organoid cultures showed that overexpression of Deltex1 results in a significant block in T-cell development and the emergence of B cells (*99*). When characterizing human early T cell progenitors, we showed that CD34^+^CD7^+^ progenitors have a specific window of Deltex1 expression (*19*), and together with the identification of CD34^-^CD10^+^CD19^+^ pre-B cells in this dataset there indeed seem to be multiple roads to intrathymic B cell development. By only including CD34^-^CD19^+^ in this dataset we likely excluded CD34^+^ B cell populations like pro-B cells, preventing characterization of an earlier developmental origin of thymic B cells. Nevertheless, using flow cytometry we previously showed that the full range of B cell developmental stages can be detected in the human thymus (*19,100*), collectively indicating the thymus as a site for human B cell development.

A central finding of this study is the thymic microenvironment’s ability to support B cell maturation, differentiation, and isotype class switching. The identification of activation-induced cytidine deaminase (AID) expression in certain subsets highlights the potential of thymic B cells to undergo antibody affinity maturation, processes traditionally associated with germinal centers (GC) in secondary lymphoid organs. Although several thymic B-cell states in our dataset express GC-associated gene programs, a physiological germinal center does not form in the healthy thymus because the structural and cellular requirements for sustained GC cycling are largely absent, including the presence of follicular helper cells (Tfh) that mature from peripheral naïve T cells, which are closely related to SP thymocytes. In an autoimmune thymic pathology setting, such as myasthenia gravis, ectopic follicle/GC structures can appear alongside strong CXCL13-driven follicular organization, highlighting that the thymus can support GC-like architecture when the niche is aberrantly created (*101*).

Against that background, the GC-like subset from our data is best interpreted as a transcriptomic state rather than evidence for a true intrathymic GC reaction. We observed a cluster with high expression of *MYC* and *CCND2* together with *CCL3/CCL4* and *NR4A1−3* expression. In secondary lymphoid organs, *MYC* is a well-established marker of strong help signals and selection in GC B cells, acting as a gatekeeper for further growth and expansion of a germinal center (*52, 53*). In our thymic context, *MYC* and *CCND2* therefore indicate that some thymic B cells temporarily engage a GC-like activation expression program, likely downstream of receptor engagement and CD40-type signals, without transitioning into a full GC reaction.

This prevention of a GC reaction is supported by our Imaging mass cytometry data, which showed the absence of true GC structures in healthy thymus. Second, the chemokines *CCL3/CCL4* expressed by GC B cells have been linked to recruitment of follicular regulatory T cells (Tfr) in second lymphoid organs, providing a built-in brake on excessive GC activity (*55*). In the thymus, a similar role is possible: chemokine production by GC-like thymic B cells could help recruit CCR5+ regulatory T-cells to prevent further GC reactions. Third, the expression of NR4A genes (encoding nuclear receptor proteins involved in apoptosis) alongside GC-like activation markers suggests that these cells have received strong BCR signaling that induces apoptosis, but are simultaneously upregulating important antagonistic programs, consistent with the tight control needed to prevent full germinal center formation in the thymus (*102*).

Without a classical GC reaction, naïve B cells still are able to differentiate into memory B cell subsets in a way that resembles an extrafollicular program (*103, 104*). At the molecular level, several B cell subsets in our data express AICDA (AID) and the BCR data in combination with the spectral flow data confirms the presence of switched isotypes across memory and plasma cell compartments. At the same time, the overall somatic hypermutation (SHM) rates remain low across subsets (∼0-1.2%). This combination, AID/CSR with limited SHM and no organized GC niche fits well with literature showing that key GC-associated processes can be initiated outside germinal centers, including class-switch recombination and even some SHM (*103, 104*). Together, these data indicate that extensive SHM is not a general feature of thymic B-cell differentiation. Rather, the accumulation of substantial IGH mutations is largely restricted to the CD27^+^ memory compartment, whereas the dominant CD21^−^memory-like and plasma cell compartments remain largely minimally mutated despite evidence of class switching elsewhere in the repertoire. This pattern is consistent with the idea that the CD27^+^ memory B cells represent a more antigen-experienced population imported from the periphery, while the major thymic CD21^−^memory-like compartment follows a differentiation route characterized by limited SHM. This is also described in a study on extrafollicular GC-independent B cell responses in lupus kidney, in which atypical B cells and antibody-secreting cells also show low rates of somatic hypermutations compared to peripheral cells (105). The CD27^+^ lgM^+^lgD^+^ population from our data resembles marginal zone memory B cells and did not show clonal overlap with the other B cell subsets, which might suggest that these cells did not originate in the thymus. Together with the fact that these cells have specialized in mounting rapid immune responses, particularly to T-independent antigens (*106,107*), it seems that these memory B cells are a subset distinct from the other thymic B cells subsets. Interestingly, CD27^+^ lgM^+^lgD^+^ B cells have also been detected in human breast milk where it was hypothesized that in newborns these cells may be involved to compensate for the low antigen-presenting capacity of newborn macrophages (*108*). The detection of these cells in the thymus of newborns would mean that after ingestion they migrate from the gut to the thymus, and interestingly these cells are the only population with the important thymus homing molecule CCR9 expression.

Most memory-like cells in our thymus dataset are CD21^−^CD27^−^B cells that resemble atypical memory B cells and carry a strong antigen-presentation profile, including expression of FCRL4/FCRL5 together with increased co-stimulatory and antigen-processing genes (CD80/CD86 and IFI30/GILT). This differs from secondary lymphoid organs, which are dominated by the classical CD27^+^ memory pool (*62*). In lymph nodes and spleen, similar CD21^low^ atypical memory-like cells are linked to extrafollicular differentiation, where B cells can become activated and generate memory- or plasmablast-like outputs without entering a stable germinal center program, particularly in settings of strong or prolonged stimulation (*103,109,110*). Our BCR data supports this interpretation. Overall, thymic B cells show high clonal diversity and little evidence for the large clonal expansions that would be expected from a strong, sustained germinal center reaction. At the same time, we do detect clear clonal overlap between specific states, most notably between an activated naïve-like cluster and the GC-like/early memory compartment, suggesting that at least a fraction of thymic B cells differentiate from an activation to memory B cell locally.

Thymic B cells are thought to acquire a specialized tolerogenic APC program through intrathymic “licensing,” in which CD40 engagement drives heightened antigen presentation and co-stimulation and, in a subset, induction of AIRE-associated transcriptional programs (*7, 33*). In this context, the rare AIRE^+^/EBI3^+^/CCL17^+^/CCL22^+^ thymic B-cell population we identify may represent a highly activated APC state that both promotes productive interactions in the medulla and expands the spectrum of self-antigens available for negative selection and Treg induction. While our data do not directly prove TRA presentation, the confinement of AIRE expression to this subset together with its co-stimulatory and chemokine profile is consistent with a role in reinforcing central tolerance alongside mTECs and dendritic cells.

Studies show that there is reason to believe that thymic B cells play a role in preventing autoimmunity (*6, 8,12, 16, 32*), but also in generating tolerance against dietary antigens (17). A recent study described repertoire relatedness between IgA1 and IgE in gastrointestinal tissue from patients with peanut allergy, suggesting that IgA1 to IgE switching could take place in the gut (*111*). With the notion that plasma cells reactive to dietary antigens have been detected in the human thymus (*17, 18*), although it was commonly believed that these antibodies are generated in lymphoid tissue in the gut (*112*). Together with the characterization of CD23^+^IGHE^+^ cells in our dataset, that in other studies has been associated with food allergy (*89*), it could well be that other dietary-specific plasma cells could also differentiate in the thymus.

A critical component of central tolerance extends beyond preventing reactivity to somatic tissues; it must also encompass tolerance to immune cell–derived antigens. In particular, developing T cells must be rendered tolerant to B cell–specific proteins. Failure to do so would predispose the host to autoreactive responses targeting B cells themselves (resulting in B cell aplasia) or immunoglobulins (manifesting as anti-globulin responses).

One plausible mechanism is that B cells load peptides derived from their own BCR and other B-cell proteins very efficiently onto MHC-II, and that this endogenous presentation is further enhanced by B-cell activation states (*113,114*). In this context, the active class-switching of memory B cell subsets we identified likely reflects a tolerance mechanism rather than a conventional humoral response observed in secondary lymphoid organs. By generating class-switched immunoglobulins in the thymus, thymic B cells can present peptides from the constant regions of IgG and IgA (and potentially primed IgE-associated programs). This process may serve to pre-emptively expose developing T cells to the diversity of the antibody repertoire, reinforcing central tolerance prior to their egress into the periphery.

The findings described in this paper provide a better understanding of the dynamics of B cell development, clonal expansion, migration, and differentiation in the human thymus, emphasizing the interconnectedness of specific subsets. Collectively, our data indicate that in addition to its primary lymphoid function, the human thymus provides a microenvironment capable of supporting secondary lymphoid functions.

## MATERIAL AND METHODS

### Isolation of cells from thymus

Thymic tissues were obtained as surgical tissue discards from children 3 to 36 months of age (median: 6 months of age) who underwent cardiac surgery; written informed consent was obtained from the parents. The children did not have immunological abnormalities. Thymocytes were isolated from the tissues by cutting the thymic lobes into small pieces and then squeezing the pieces through a metal mesh; the cells were then frozen and stored in liquid nitrogen until further use.

### Flow cytometry and cell sorting

Spectral flow cytometry and cell sorting were performed at the Leiden University Medical Center Flow Cytometry Core Facility using a Cytek Aurora 3L (Cytek Biosciences, Fremont, CA) and a BD FACS Aria III 4L (BD Biosciences, San Jose, CA), respectively. Non-T lineage cells were sorted using antibodies against the following surface markers: CD16, CD56, CD13, CD33, CD19, and CD34. The purity of the sorted populations is shown in **Fig. 1A**.

### Single-cell RNA sequencing

To match the experimental conditions of our previously generated dataset, cell suspensions from alternative lineage and CD34^+^ were barcoded (10x Chromium Single Cell platform, 10X Genomics) using the Chromium Single Cell 5’ Library (10x Genomic) and used to generate two single-cell 5⍰ gene expression libraries using the 10x version 1.1 chemistry for 5’ sequencing. The loaded cell numbers ranged from 300 to 500,000, aiming for 5000 cells per reaction. All libraries were sequenced using an Illumina NovaSeq 6000 (paired-end 75-bp reads) to an average depth of 50,000 reads per cell, resulting in two datasets.

### Preprocessing of scRNA-seq data

Sequencing data from the Alternative lineage dataset was demultiplexed and mapped against the GRCh38 genome using Cell Ranger 7.1.0. The resulting gene count matrices were loaded into R using Seurat v4. We removed all genes related to TCR and BCR from the original count matrix to prevent clustering biased towards the TCR or BCR repertoire in further downstream analyses. Seurat objects were created including cells with >100 genes expressed and excluding genes expressed in fewer than 3 cells. We used the PercentageFeatureSet function to calculate the percentage of mitochondrial gene expression per cell. Before clustering cells with >30% mitochondrial reads were removed.

We used the Vireo method based on genotyping to identify the donors *in silico* for every cell and used the genotyping profiles to detect doublets. In addition to doublets detected by genotype we used scDblFinder (https://github.com/plger/scDblFinder) to detect additional doublets in our data. We used the identified doublets for model training for scDblFinder by providing the barcodes of genotyped detected doublet cells using the knownDoublets and knownUse arguments. We set the expected doublet rate, i.e. the proportion of the cells expected to be doublets to 0.008, (0.8%) based on the doublet rates 10x Genomics expects for 5’ sequencing libraries. Detected doublets were subsequently removed from data before removing low-quality cells which we identified using a cluster-based approach: after Normalization and scaling we performed PCA analysis using the 2000 most variable genes. After PCA analysis and clustering analysis with the FindNeighbours (dims 1:50) and FindClusters function (resolution = 1), 40 principal components were used for UMAP generation. Clusters representing low-quality cells, such as those with high mitochondrial gene content, low total RNA counts, or low unique feature counts, were identified and removed. The criteria for low-quality clusters were determined based on their distinct transcriptional profiles and/or their metadata attributes, such as high percentages of mitochondrial gene expression and low percentages of ribosomal gene expression.

Based on these criteria we removed 15 clusters from our dataset. Following the removal of these clusters, the remaining high-quality cells were retained for downstream analyses. Finally, we removed cells without a genotype in order to correct the data for donor effects, which resulted in a dataset consisting of 4796 cells.

After filtration, we split the dataset into 3 subsets based on the 3 identified genotypes. We used CCA in Seurat to identify common sources of variation between the donors. After identifying anchors using 30 principal components, the dataset was integrated, and the integrated dataset was used for dimensionality reduction. Before running the PCA dimensionality reduction, cell cycle scores were calculated and used for cell cycle regression based on the expression of genes involved in the cell cycle. The residuals were used to scale the dataset. After PCA analysis clustering analysis was performed with the FindNeighbours (dims 1:50) and Findclusters function (resolution = 1) which identified 14 clusters. We then visualized cells using a twodimensional UMAP plot generated using the RunUMAP function in Seurat package with 25 principal components. After scaling the RNA data, the FindAllMarkers function in Seurat was used to identify differentially expressed genes between the 14 clusters which were used for cell type annotation. After clustering we included the counts for TCR and BCR genes to the Seurat object.

### Demultiplexing cells to donors

To determine the identity of the individual donor of every cell, we used the Bayesian demultiplexing tool Vireo (v 0.4.2, R version). In brief, we first generated a list of single nucleotide polymorphism (SNP) positions by aligning all expressed reads from all cells and selecting positions with a minimum allele frequency of 0.1 and minimum total coverage of 20. Next, in each cell and at each position we identified overlapping SNPs and counted them in two disjoint groups corresponding to the reference and non-reference alleles. The allelic count matrices were then used to fit a Vireo model that either identified the most likely donor for each cell or classified the cell as a doublet.

### Cell Type Annotation

Cell type annotation was performed using several approaches:

1. **Manual:** The FindAllMarkers function in Seurat was used to identify differentially expressed genes between the 14 clusters and literature reviews were conducted to annotate clusters.
2. **Automated:** Using Seurat’s Azimuth tool in Seurat version 5, a reference-based annotation was performed. Processed single-cell RNA sequencing data were separately integrated with reference atlases of human PBMC, human bone marrow cells and two human tonsil cell atlases (*23, 24*) using the Azimuth pipeline. This integration involved the transfer of cell type labels from the reference dataset to the query dataset through canonical correlation analysis (CCA) and k-nearest neighbor (kNN) mapping. To refine these annotations, manual curation was performed using the cluster marker genes identified by Seurat to align with biologically relevant markers reported in published studies (*23, 24*).

The final cell type annotations represent a combination of automated labeling from Azimuth and manual verification using validated marker genes. One small cluster was identified as epithelial cells, based on high expression of CCL19 and lack of B makers (CD19, MS4A1) and was removed from further analyses.

### Single-cell BCR repertoire analysis

Raw reads from the gene expression sequencing runs were processed using cellranger_vdj in Cell Ranger (v.7.1.0) with a custom reference provided by the manufacturer (version 2.0.0 GRCh38 VDJ-alts-ensembl). We separately ran cellranger_vdj for BCR reconstruction using the -chain=IG argument.

BCR contigs contained in all_contigs.fasta and all_contig_annotations.csv were then processed further using dandelion singularity container (v.0.2.4) (https://www.github.com/zktuong/dandelion). BCRs were then matched to cell barcodes with dandelion (115). Dandelion identified BCR rearrangements in 4547 cells, with complete BCR rearrangements in 2665 cells which were used for isotype classification.

BCR sequences were reconstructed directly from 5’ single-cell gene expression sequencing data using the Cell Ranger VDJ pipeline (10x Genomics, v7), without a separate VDJ-enriched library. Cell Ranger output (all_contig_annotations.csv) was re-annotated using Dandelion to obtain IMGT-aligned germline sequences (*116*). Contigs were filtered for productive rearrangements with ≥2 UMI counts. V-region mutation frequencies were calculated using SHAzaM (observedMutations) (*117*) with IMGT_V_BY_REGIONS region definitions, providing separate replacement (R) and silent (S) mutation frequencies for CDR1, CDR2, FR1, FR2, and FR3 regions. Aggregate CDR (CDR1+CDR2) and FR (FR1+FR2+FR3) mutation frequencies were computed per contig. R/S ratios were calculated per region to assess selection signatures, with a ratio of ∼2.9 representing the random expectation. The fraction of somatically mutated cells was defined using a mu_freq threshold of >0.01.

Clonal analysis was performed using scRepertoire2 (*118*). The combineBCR() function was used to pair singlecell barcodes with corresponding heavy and light chain sequences, with analyses performed separately for all donors combined and per individual donor. Clonotypes were defined using the “strict” call (CDR3 amino acid sequence + V-gene usage for both paired chains). The percentage of unique clonotypes per cluster was calculated using clonalQuant(), and clonal size distributions were assessed using clonalAbundance(), both analyzed separately for IGH, IGL/IGK, and paired chains. Clonal diversity was evaluated using clonalDiversity(), which computes Shannon entropy, inverse Simpson’s index, normalized entropy, and Gini-Simpson index. CDR3 length distributions were analyzed using clonalLength() for both IGH and IGL/IGK chains, with amino acid length as the metric. V-gene usage was visualized using vizGenes() and percentGenes(), and V-J gene pairing frequencies were assessed using percentVJ(). All repertoire analyses were performed both across 14 B cell clusters and across four functional groups (naive B, CD21^−^memory-like B, CD27^+^ MBC, and plasma cells), and additionally by isotype class (lgM/lgD, lgG, lgA).

Clonal overlap between clusters was assessed using clonalOverlap() with overlap coefficient, Morisita index, Jaccard index, cosine similarity, and raw count methods, applied both across all donors and per individual donor. To identify clonally related cells, IGH contigs were deduplicated to one per cell (highest UMI) and grouped into clones within each donor based on shared V-gene and J-gene usage (allele-stripped) and CDR3 amino acid similarity ≥80% (normalized Levenshtein distance, single-linkage hierarchical clustering). Interdonorclone sharing was assessed and excluded. Clonal groups containing both class-switched (IgG/lgA/lgE) and unswitched (IgM/IgD) members within the same donor were identified as putative intrathymic class-switch recombination (CSR) events. The clonal expansion rate was defined as the percentage of cells belonging to clones with ≥2 members. For IGL/IGK, clones were defined by exact CDR3 nucleotide identity combined with V-gene and J-gene usage within donor and chain, to avoid false matches from convergent light chain recombination.

### Tonsil reference atlas analysis

To compare thymic memory B cell phenotypes with their secondary lymphoid organ counterparts, we reanalyzed the publicly available human tonsil cell atlas (24). The annotated Seurat object containing naive and memory B cells (20230911_NBC_MBC_seurat_obj.rds) was obtained from Zenodo (119). To match the age range of our pediatric thymic cohort, only donors aged 3, 4 and 5 years were retained for analysis. Cluster annotations as provided by the original authors (annotation_20230508) were retained throughout. Memory B cell (MBC) clusters were defined as Early MBC, ncsMBC, ncsMBC FCRL4/5^+^, csMBC, csMBC FCRL4/5^+^, and MBC FCRL5^+^. Expression of *CD27* and *AICDA* across these clusters was compared between the different memory B cell clusters. To benchmark CD27 expression in tonsil against the CD27^+^ and CD27^−^MBC clusters identified in our thymic dataset, the percentage of CD27^+^ versus CD27^−^cells across all tonsil MBC clusters was computed per donor and visualized as barplots showing mean ± standard error with individual donor values overlaid.

### Thymus tissue preparation for Mass Cytometry

Fresh thymic lobes were sectioned into small pieces (1−2 mm thick), embedded in optimal cutting temperature (OCT) compound, snap-frozen in isopentane, and stored at −80°C until further processing. Tissue sections were prepared as previously described (*120*). Briefly, 5−μm cryosections were cut onto silane−coated glass slides (VWR), air−dried for 1 hour at room temperature (RT), and fixed in 4% paraformaldehyde (PFA) for 10 minutes. The slides were then washed and rehydrated in DPBS before being blocked with SuperBlock (Thermo Fisher) at RT for 30 minutes. Following blocking, slides were incubated overnight at 4°C with the following antibody mix: CD45−89Y (clone H130) − 1:50; CD27−167Er (clone 0323) − 1:50; CD38−172Yb (clone HIT2) − 1:100; CD3−170Er (clone UCHT1) − 1:100; CD20−161Dy (clone H1) − 1:50; COL1−147Sm (polyclonal) − 1:100; HLA−DR−168Er (clone L243) − 1:800; CD8a−146Nd (clone RPA−T8) − 1:50; CD4−145Nd (clone RPA−T4) − 1:50). After incubation, slides were washed and stained with the DNA intercalator Ir (1:400 dilution) for 30 minutes at RT. Finally, slides were rinsed with DPBS followed by Milli-Q water and dried using compressed air.

### Mass Cytometry Image acquisition

Tissue samples were processed using a Helios time−of-flight mass cytometer integrated with the Hyperion Imaging System (Fluidigm). All imaging mass cytometry (IMC) procedures followed the standard protocol provided by Fluidigm. In summary, after purging the ablation chamber with helium, tissues were subjected to ultraviolet (UV) laser ablation in a spot-by-spot manner, operating at a resolution of 1 μm and a frequency of 200 Hz. Areas of interest measuring 1,000 μm x 1,000 μm were selected, with 5 to 8 such regions analyzed per tissue section. Marker intensity was visualized through the Fluidigm MCD™ Viewer (vl.0.560.2) and threshold minimum values were adjusted manually foreach marker using the MCD™ Viewer.

### Spectral flow cytometry of thymic B-cell subsets

Single-cell suspensions were prepared from two human thymus samples (overlapping with two scRNA-seq samples) and stained with a custom surface-only spectral flow cytometry panel for phenotypic analysis of thymic B-cell subsets and surface immunoglobulin isotypes. The antibody panel included CD27-BV421 (BD), kappa-BV480 (BD), IgM-BV510 (BioLegend), CD45RA-BV570(BioLegend), CD23-BV605 (replacing CD62L in the original panel), CD24-BV650 (BD), CD21-BV711 (BD), lambda-BV750 (BD), CD19-BV786 (BD), IgD-FITC (BioLegend), CD20-PE-CF594 (BD), CD45-Alexa Fluor 700 (BD), and CD38-APC-H7 (BD). The panel further contained surface immunoglobulin reagents forlgGl, lgG2, lgG3, lgG4, IgAl, and lgA2 (Cytognos). No intracellular immunoglobulin staining was performed. Samples were acquired on a spectral flow cytometer (Cytek Aurora) and analyzed in Infinicyt (Cytognos).

For analysis, events were first gated on lymphocyte size and granularity using forward and side scatter, followed by exclusion of doublets by singlet gating. B cells were identified as CD19^+^CD20^+^ cells. Within the B-cell compartment, cells were separated into pre-GC B cells and memory B cells using CD27 and surface IgM, as shown in the representative gating strategy in Supplementary Figure 1. Pre-GC/naïve B cells were defined as CD27^−^lgM^+^cells lacking class-switched surface immunoglobulin expression and were further subdivided according to CD21 expression into CD21^+^and CD21^−^fractions. Naïve B-cell identity was confirmed by coexpression of surface IgM and IgD.

Memory B cells were defined as cells that were CD27^+^ and/or class-switched on the basis of surface immunoglobulin staining. Within this compartment, the Infinicyt analysis hierarchy resolved non-switched lgM^+^lgD^+^ memory B cells, lgGl^+^, IgG2^+^, IgG3^+^, IgG4^+^, lgAl^+^, and IgA2^+^ memory B cells, and IgD-only memory cells. The non-switched memory compartment was analyzed within the CD27^+^gate, whereas class-switched memory populations were further separated into CD27^+^and CD27^−^ fractions. These memory subsets were subsequently subdivided according to CD21 and CD23 expression, allowing discrimination of CD21^+^CD23^+^, CD21^+^CD23^−^, CD21^−^CD23^+^, and CD21^−^ CD23^−^ populations. Surface isotype usage within these gated compartments was then assessed using the immunoglobulin isotype panel. Population frequencies were calculated in Infinicyt as percentages of total CD19^+^CD20^+^ B cells or of the relevant parent gate, as indicated in the figure panels. Event counts per gated population were exported and analyzed in R using ggpubr.

### Statistical Analysis

Statistical analyses were conducted using the comparejneans function from ggpubr in R. P-values were calculated with the Wilcoxon test adjusted for multiple testing using the Benjamini-Hochberg correction. A p-value < 0.05 was considered statistically significant.

## Data Visualization

We used BBrowserS (version 2.9.23) (*121*) to visualize the Seurat objects and to create UMAP plots, gene expression, gene signature plots, pie charts, TRBV-J heatmaps and donor proportion plots. Dotplots and Violinplots were created using the Dotplot() and VlnPlot() functions in Seurat. Clonotype analysis visualizations were created using the plot functions within scRepertoire2. Additional boxplots, proportion plots and density plots were generated using ggplot2 in R.

## Acknowledgements

We are grateful to the Leiden University Medical Center Flow Cytometry Core Facility for assistance with cell sorting. K.C.-B. and F.J.T.S. are supported in part by Novo Nordisk Foundation grants (NNF21CC0073729) that support the Novo Nordisk Foundation Center for Stem Cell Medicine at LUMC.

## Supplementary Figure Legends

**Supplementary Figure 1.**
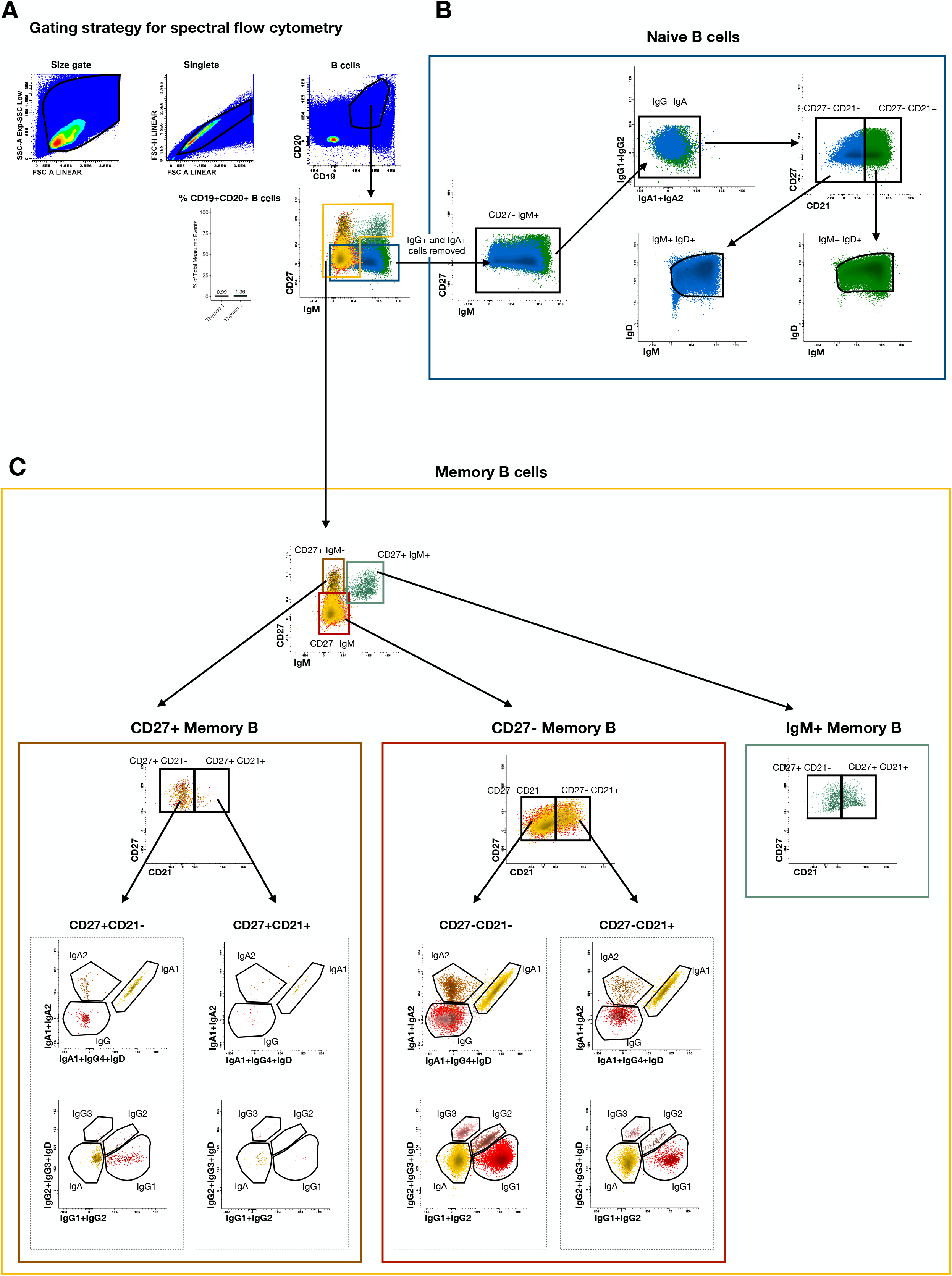
Gating strategy for spectral flow cytometric analysis of thymic B-cell subsets. Events were first gated on lymphocyte size and granularity, followed by exclusion of doublets to obtain singlets. B cells were identified as CD19^+^CD20^+^ cells. Within the B-cell compartment, naïve B cells (blue box) were defined as CD27^−^cells lacking switched isotypes (IgG and IgA) and were further subdivided into CD21^+^ and CD21^−^fractions, both showing an lgM^+^lgD^+^phenotype. Memory B cells were defined as CD27^+^ and/or class-switched cells and separated into CD27^+^ memory B cells, CD27^−^ class-switched memory-like B cells, and IgM^+^ memory B cells. The CD27^+^ and CD27^−^ class-switched memory compartments were further subdivided based on CD21 expression and analyzed for surface isotype usage, including IgG subclasses and lgA1/lgA2. Representative plots are shown for each gating step.

**Supplementary Figure 2.**
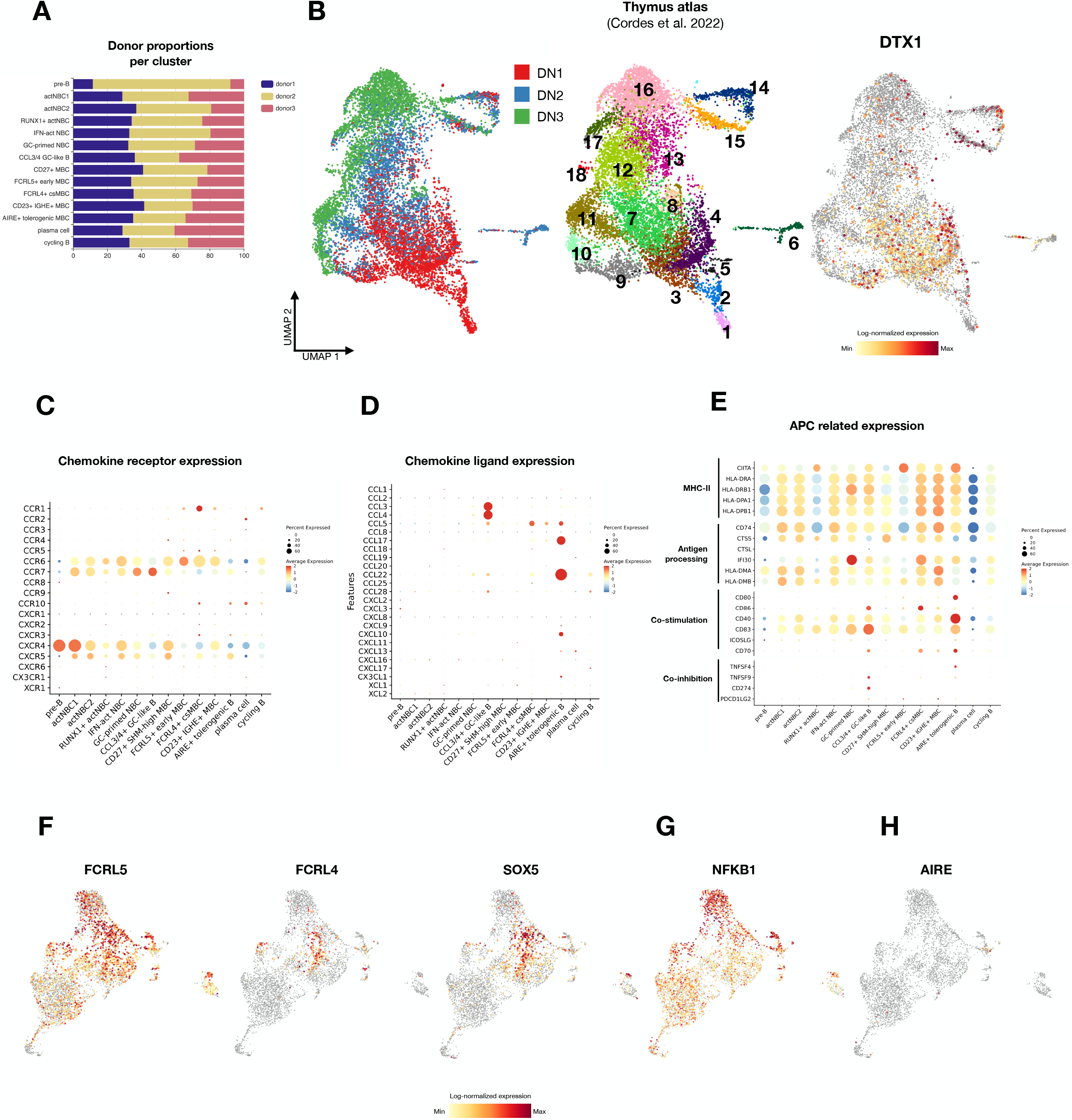
Extended characterization of thymic memory-like B-cell subsets with APC-like features. **(A)** Relative contribution of each donor to the annotated thymic B-cell clusters. **(AB)** Projection of DN1, DN2, and DN3 subsets from Cordes et al., 2022, highlighting expression of DTX1 in thymocytes during early T cell development. **(C)** Dot plot showing expression of chemokine receptor genes across thymic B-cell clusters. **(D)** Dot plot showing expression of chemokine ligand genes across thymic B-cell clusters. **(E)** Dot plot depicting expression of genes associated with MHC class II, antigen processing, co-stimulation, and co-inhibition. **(F)** UMAP expression plots of FCRL5, FCRL4, and SOX5. **(G)** UMAP expression plot of NFKB1 expression. **(H)** UMAP expression plot of AIRE expression, restricted to a rare thymic B-cell subset.

**Supplementary Figure 3.**
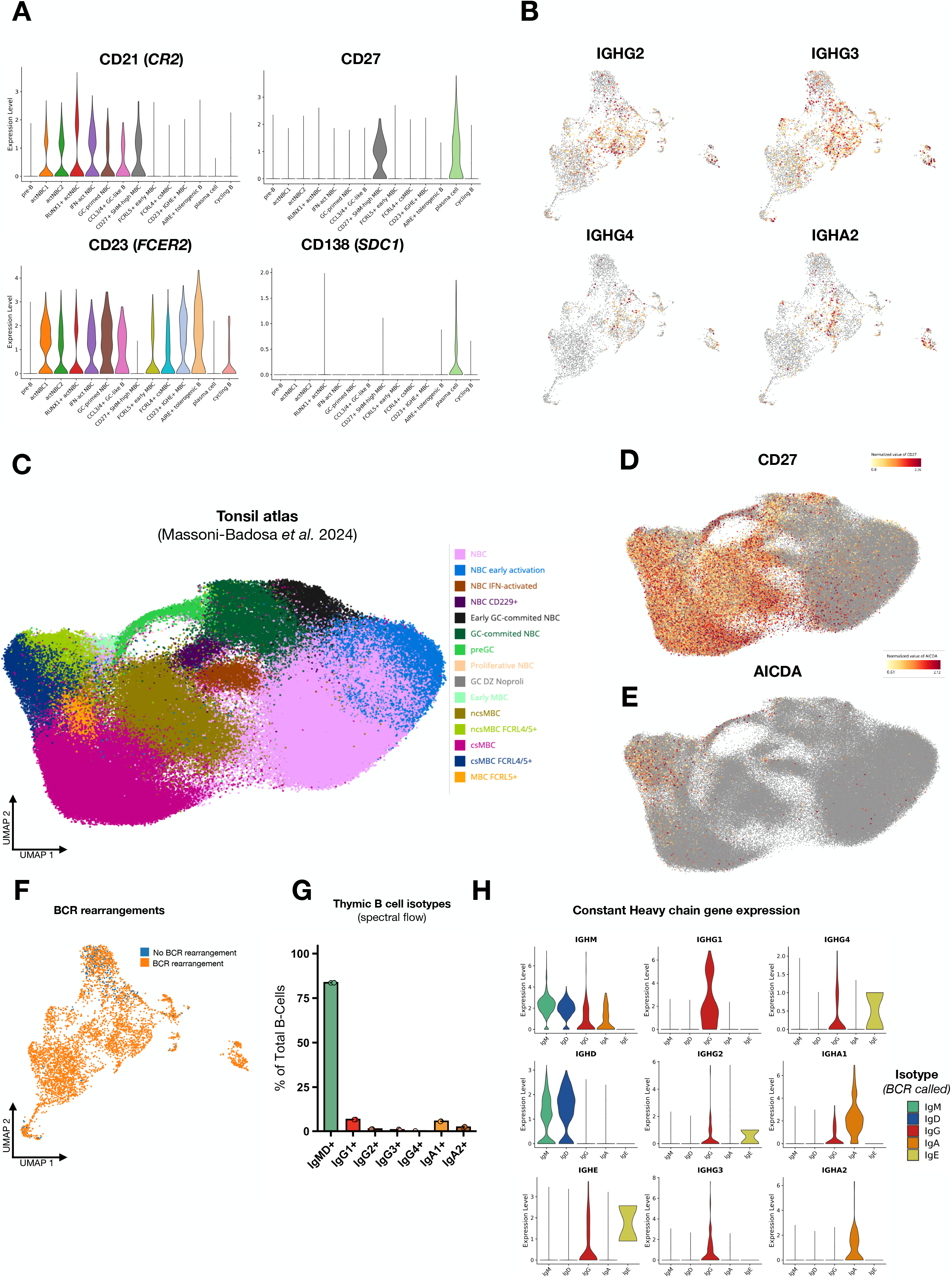
Additional marker expression and isotype characterization of thymic B-cell subsets. **(A)** Violin plots showing expression of selected surface marker genes across thymic B-cell clusters, including CD21 (CR2), CD23 (FCER2), CD27, and CD138 (SDC1). **(B)** UMAP feature plots showing expression of additional immunoglobulin heavy-chain constant region genes IGHG2, IGHG3, IGHG4, and IGHA2. **(C)** Annotated UMAP of a published human tonsil B-cell atlas (Massoni-Badosa et al., 2024), shown for comparison with thymic B-cells. **(D)** UMAP feature plot of CD27 expression in the tonsil dataset. **(E)** UMAP feature plot of AICDA expression in the tonsil dataset. **(F)** UMAP showing the distribution of cells with and without successfully reconstructed BCR rearrangements in the thymic B-cell dataset. **(G)** Surface isotype composition of total B cells measured by spectral flow cytometry. **(H)** Violin plots comparing BCR-assigned isotype class with expression of the corresponding heavy-chain constant region genes, including IGHM, IGHD, IGHG1–4, IGHA1-2, and IGHE.

**Supplementary Figure 4.**
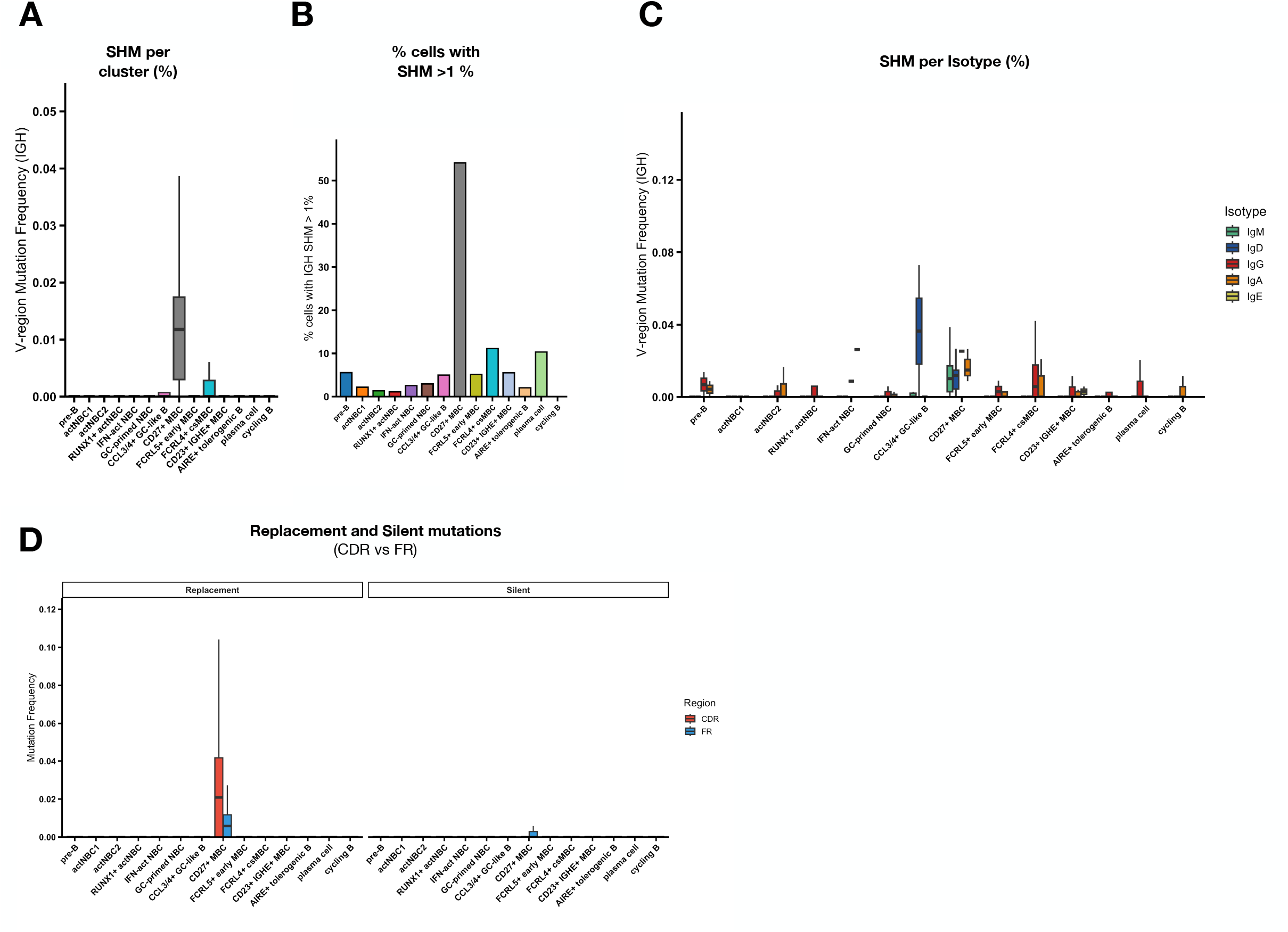
Cluster-level overview of IGH somatic hypermutation across thymic B-cell subsets. **(A)** Distribution of IGH V-region mutation frequencies across all annotated thymic B-cell clusters. SHM frequencies were calculated as the number of nucleotide mutations in the IGH V region divided by the total number of nucleotides analyzed. **(B)** Fraction of cells in each cluster with >1% IGH SHM, shown to identify clusters containing a substantial proportion of clearly mutated sequences. **(C)** IGH SHM frequencies displayed by BCR-assigned isotype for each thymic B-cell cluster. **(D)** Distribution of replacement and silent mutations in complementarity-determining regions (CDR) and framework regions (FR) across all thymic B-cell clusters.

**Supplementary Figure 5.**
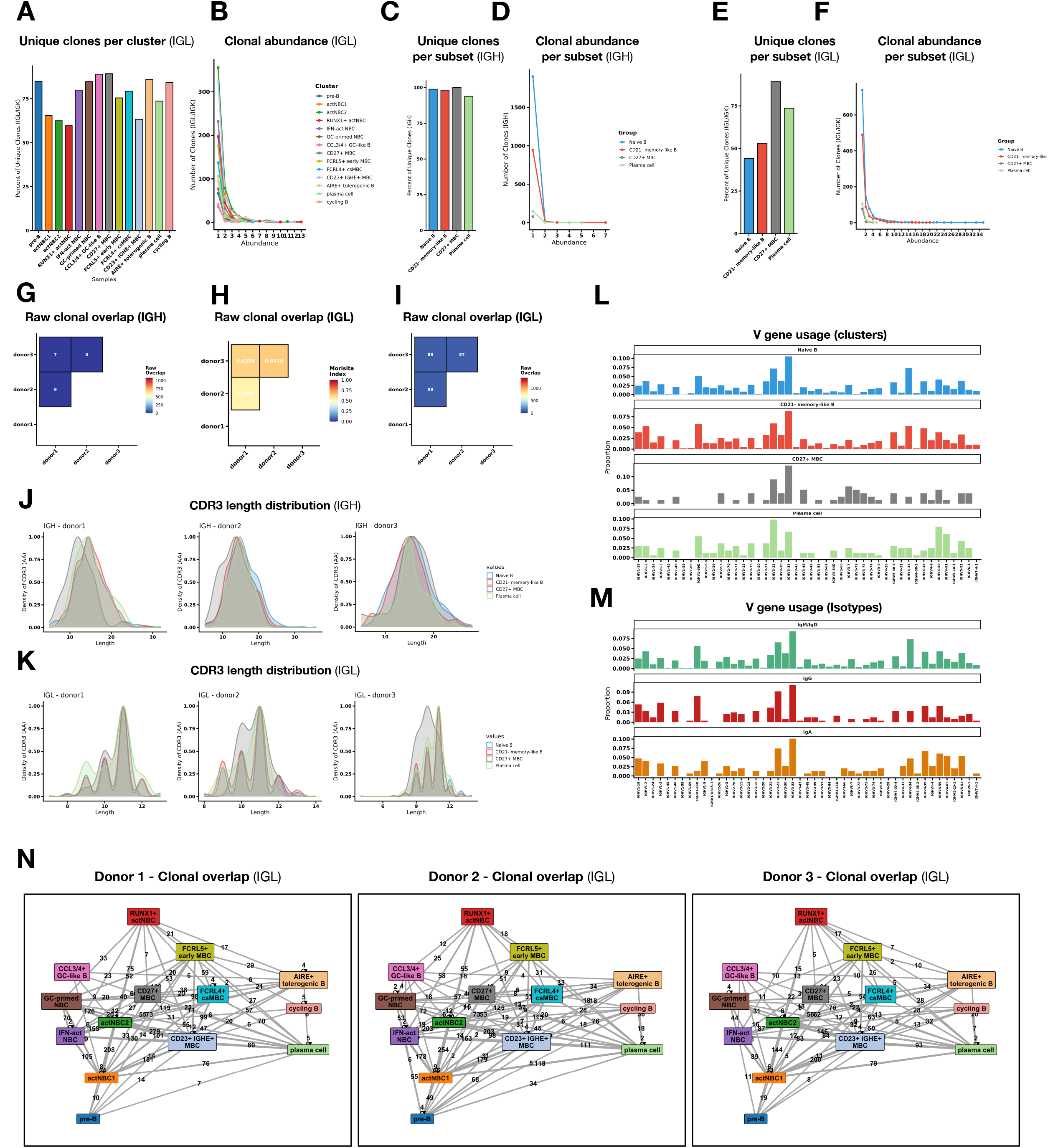
Additional repertoire features of thymic B-cell subsets based on IGH and IGL analyses. **(A)** Proportion of unique IGL clones within each annotated thymic B-cell cluster. **(B)** Relative abundance (number of occurrences of clonotypes) distribution of IGL-defined clonotypes across thymic B-cell clusters. **(C)** Proportion of unique IGH clones within the four major thymic B-cell compartments (Naïve B, CD21^−^memorylike B, CD27^+^ MBC, and plasma cells). **(D)** Relative abundance (number of occurrences of clonotypes) distribution of IGH-defined clonotypes across the four major thymic B-cell compartments. **(E)** Proportion of unique IGL clones within the four major thymic B-cell compartments. **(F)** Relative abundance (number of occurrences of clonotypes) distribution of IGL-defined clonotypes across the four major thymic B-cell compartments. **(G)** Absolute numbers of clones overlapping donor-to-donor based on **IGH** repertoires, calculated using the Morisita–Horn index. **(H)** Donor-to-donor clonal overlap based on IGL repertoires, calculated using the Morisita–Horn index, showing minimal overlap between donors. **(I)** Absolute numbers of clones overlapping donor-to-donor on **IGL** repertoires. **(J)** Density plots of IGH CDR3 amino-acid length distributions for each individual donor across the four major thymic B-cell compartments. **(K)** Density plots of IGL CDR3 amino-acid length distributions for each individual donor across the four major thymic B-cell compartments. **(L)** IGH V-gene usage across the four major thymic B-cell compartments. **(M)** IGH V-gene usage by isotype. **(N)** Within-donor IGL clonal overlap networks for each donor, showing shared clone groups between thymic B-cell clusters. Edge thickness reflects the number of overlapping clones.

## Notes

### Competing Interest Statement

The authors have declared no competing interest.

